# Sample size buys detection, not localisation: an identifiability limit for hippocampal subfield morphometry

**DOI:** 10.64898/2026.08.19.745596

**Authors:** Rodrigo Debona, Roger Walz

## Abstract

Automated segmentation has made hippocampal subfield volumes a routine measurement, and studies now report which subfield relates to an outcome rather than whether the hippocampus does. Those reports do not agree with one another, and the standard explanation is insufficient statistical power. We argue that a second limit operates independently of sample size. Using 638 participants from a population-derived adult lifespan cohort, we first show that no individual subfield contributes to a general cognitive factor beyond a single global size component: no coefficient interval excludes zero, the local block carries half a percent of outcome variance, and no model improves out-of-sample prediction over the global factor alone. Because an observed null cannot distinguish an absent effect from an effect the design cannot locate, we then planted effects of known location and size in the measured design and in a whitened copy of it that preserves sample size, dimensionality and effect size while removing only the correlation between subfields. The arms were paired down to the noise vector. Collinearity did not place recovery out of reach; it multiplied the required sample size by a factor of roughly two to three, and the penalty widened as cohorts grew. At the effect sizes this literature reports, neither design reached an adequate recovery rate at any sample size, and coarsening the parcellation rescued neither. The choice of estimator moved recovery further than collinearity did. We provide a calibration surface on which a planned design can be located before data collection.

## 1 Introduction

Automated segmentation made the internal structure of the hippocampus a routine measurement. Since the probabilistic atlas of Iglesias et al. (2015) was released with FreeSurfer, a standard-resolution structural scan yields volumes for a dozen or more subfields, and studies of ageing, memory and disease now report which of them relates to an outcome rather than whether the hippocampus does. The measurement is cheap, the labels are anatomically named, and the resulting claims are correspondingly specific.

The claims do not agree with one another. Two decades ago, Van Petten (2004) synthesised 33 studies of whole hippocampal volume and memory and concluded that the association probably does not exist among young adults, emerging only in samples weighted toward older participants. A more recent meta-analysis in typically developing children and adolescents reported the opposite direction (Botdorf et ah, 2022). At the level of subfields the picture is no steadier: individual reports nominate different structures, and the nominations do not obviously track sample size, cohort or acquisition. When quantitative syntheses of the same construct disagree about the sign of an effect, the explanation is unlikely to lie in the design of any single study.

The prevailing diagnosis for inconsistency in brain-behaviour research is statistical power. Marek et al. (2022) showed that the largest replicable brain-wide associations are small, reaching r = 0.14 for univariate and r = 0.34 for multivariate methods, and argued that reproducibility therefore requires thousands of participants. The claim has been contested, with multivariate models shown to replicate at moderate sample sizes under favourable conditions (Spisak et ah, 2023) and a reply from the original authors defending the original estimates (Tervo-Clemmens et ah, 2023); a parallel argument holds that measurement design can substitute for enrolment (Rosenberg and Finn, 2022; Gratton et ah, 2022). What every position in that debate has in common is its object. All of them concern whether an association reproduces. None asks whether the anatomical location of that association is recoverable, and the two questions have no reason to share an answer.

There is a reason to expect they do not. Recovering which predictors carry an effect is a different problem from establishing that some do, and it imposes conditions on the design itself. For l_1_ methods the requirement is an irrepresentable condition, which fails when the predictors carrying the effect are strongly correlated with those that do not (Zhao and Yu, 2006), and recovery thresholds scale as a design-dependent constant multiplying k log(p—k), with that constant determined by the covariance of the predictors (Wainwright, 2009). Hippocampal subfields are an unusually unfavourable case. They are anatomically adjacent, they are parts of a whole and therefore constrained to sum, and their boundaries are placed by a shared prior at a resolution where several of those boundaries are not visible in the image. Their correlation belongs to the measurement, not to the sample, so enrolling more participants does not dilute it. The genetics literature has effectively conceded the point in practice: the largest genome-wide analysis of subfield volumes co-varied for whole hippocampal volume precisely in order to recover signal specific to individual subfields (van der Meer et ah, 2020), a step the cognitive literature never standardised.

What has not been done is to measure what a real subfield design can recover. Reliability studies quantify measurement error (Brown et ah, 2020; Chiappiniello et ah, 2021), agreement studies quantify divergence between pipelines (Samara et ah, 2021), and power analyses quantify detection, but none of these states the probability that an analysis will name the correct subfield when a subfield effect genuinely exists. That probability cannot be estimated from observational data alone, because an observed null is compatible with two incompatible explanations: that no effect is present, or that an effect is present and the design cannot locate it. The two call for opposite responses from the field, and distinguishing them requires knowing what the design would do if an effect were there.

We supply that measurement. Using 638 participants from the Cam-CAN cohort (Shafto et ah, 2014; Taylor et ah, 2017), we first characterise the observed association between 36 hippocampal subfield volumes and a general cognitive factor, separating a global size component from the local contrasts orthogonal to it and asking, with a regularized horseshoe prior (Piironen and Vehtari, 2017), whether any subfield contributes beyond that global component. We then plant effects of known location and known size in the measured design, and in a whitened copy of the same design that preserves sample size, dimensionality and effect size while removing only the correlation among subfields, and we recover them with the same model. The two arms are paired down to the noise vector, so the difference between them isolates the contribution of the geometry. To our knowledge this counterfactual, which the recovery-threshold results anticipate in theory, has not previously been constructed empirically in a neuroimaging design.

The experiment settles the question with a number. Collinearity among subfields does not place an adequate result out of reach; it raises the price of one, by a factor we estimate, and that price grows as cohorts get larger instead of shrinking. At the effect sizes this literature actually reports, though, the ceiling binds for both arms. Neither the measured design nor its orthogonalised counterpart reaches an adequate recovery rate at any sample size, and coarsening the parcellation rescues neither. A third result was not anticipated: the choice of estimator moves recovery further than the presence of collinearity does. That is a warning about reporting practice as much as a finding about the hippocampus. What the study delivers for use is a calibration surface on which a planned design can be located before data are collected, and a case for naming equivalence classes of indistinguishable labels in place of individual winners.

## 2 Methods

### 2.1 Participants

Data came from stage 2 of the Cambridge Centre for Ageing and Neuroscience cohort (Cam-CAN; Shafto et ah, 2014; Taylor et ah, 2017), a cross-sectional population-derived sample stratified by age decade. Of the 653 participants with hippocampal subfield segmentations, 638 had complete data on the primary outcome and all covariates and were analysed. Exclusions were driven by missing values rather than by any criterion applied after inspection of the results.

### 2.2 Hippocampal subfield volumes

Structural images were processed with FreeSurfer 8.2.0 (Fischl, 2012; Fischl et ah, 2002). Subfield volumes were produced by the hippocampal subfield module (segmentHA_Tl. sh), which implements the probabilistic atlas of Iglesias et al. (2015): an atlas built from fifteen autopsy specimens scanned at approximately 0.13 mm isotropic resolution and manually labelled into thirteen substructures, applied to in vivo images through a generative Bayesian model that adapts to contrast and resolution. The anterior and posterior subdivision of the subfields, which the software labels head and body, dates from that atlas and has been available since FreeSurfer 6.0; the module has not changed substantially across subsequent releases (Samann et ah, 2022).

Volumes were exported with asegstats2table and columns were selected from the standard output. No mask was constructed and no volume was recomputed. Columns corresponding to amygdala nuclei, to aggregate labels (whole hippocampus, hippocampal head, hippocampal body) and to the hippocampal fissure were removed bilaterally, leaving 18 labels per hemisphere and 36 in total: parasubiculum; presubiculum, subiculum, CA1, CA3, CA4, the granule cell and molecular layer of the dentate gyrus, and the molecular layer, each divided into anterior and posterior segments; fimbria; the hippocampus-amygdala transition area; and the hippocampal tail. The fissure was removed because it is a cerebrospinal fluid space rather than a subfield, and the aggregate labels because they are sums of the retained ones. CA2 is not a separate label in this atlas: the label denoted CA3 corresponds to the combined CA2 and CA3 field, and is referred to as CA3 throughout for consistency with the software output. Subsegments are referred to as anterior and posterior throughout, rather than head and body, following the convention of The Hippocampus Book; the two namings denote the same partition. Whole hippocampal volume was retained separately as the predictor of the baseline model.

Reliability across these labels is not uniform. In the release on which the published reliability work was performed, test-retest agreement is high for the molecular layer and dentate gyrus, with intraclass correlations above 0.95, and lower for the parasubiculum, with intraclass correlations between 0.78 and 0.89, mean volume differences above 5% and overlap below 70% (Brown et al., 2020); multicentre work reports comparable limitations for the fimbria (Chiappiniello et al., 2021). The parasubiculum, fimbria and hippocampus-amygdala transition area, six labels across the two hemispheres, were flagged on that basis and a sensitivity analysis excluding them was specified in advance. Both reliability studies used FreeSurfer 6.0; extending their conclusions to 8.2.0 assumes the module has not changed materially, which the ENIGMA review supports (Samann et ah, 2022) but which was not verified here.

### 2.3 Cognitive outcome

The primary outcome was CognitiveG, the first principal component of the Cam-CAN cognitive battery. The second and third components were retained as secondary outcomes for the specificity analysis. No transformation was applied beyond the standardisation described below.

### 2.4 Preprocessing and construction of the design

Volumes were log_10_ transformed, so that multiplicative head-size scaling enters additively, and winsorised at four standard deviations within each column. Both steps were fixed before any model was fitted.

Nuisance adjustment used a single design matrix containing an intercept, log_10_ estimated intracranial volume, age as a natural cubic spline with five quantile knots, sex and years of education. Estimated intracranial volume entered as a covariate rather than as a divisor, because regional volumes do not scale proportionally with intracranial volume and the proportion method is biased whenever intracranial volume is itself associated with a covariate of interest. Voevod-skava et al. (2014) show that regional volumes do not scale proportionally with intracranial volume, that the proportional method inverts the sign of the residual association with intracranial volume, and that it leaves spurious sex differences which the residual method removes. Age entered as a spline because subcortical decline across the adult lifespan is nonlinear (Fjell et ah, 2013; Bethlehem et ah, 2022). Both the subregion volumes and the outcome were residualised on this matrix, which by the Frisch-Waugh-Lovell theorem is equivalent to entering the nuisance terms unpenalised alongside the predictors of interest, and keeps the shrinkage prior away from the covariates.

The residualised volumes were standardised, and a global factor was formed as the standardised row mean of those columns. An unweighted mean was used rather than a first principal component so that the factor does not depend on the sample. Local contrasts were then obtained by residualising each standardised column on the global factor and restandardising, which makes every local column exactly orthogonal to the global factor while leaving the correlation among the local columns untouched. That residual correlation is the object of the study and was not removed.

### 2.5 Models

Four models were fitted to the observed data. M—1 regressed the outcome on nuisance-adjusted whole hippocampal volume alone. M0 replaced it with the global factor. M1, the model reported throughout, entered the global factor with a weakly informative prior and the 36 local contrasts under a regularized horseshoe prior (Piironen and Vehtari, 2017), with the prior guess for the number of relevant predictors set to four, a slab of four degrees of freedom and unit scale. M2 applied the horseshoe to all standardised volumes without the global-local split. Two further models were fitted as sensitivity analyses: M1 restricted to the labels without documented reliability limitations, and a compositional version in which centred log-ratio coordinates replaced the local contrasts and the log geometric mean size replaced the global factor. A multivariate model over the three cognitive components estimated outcome specificity.

Observation models were Student-t with an estimated degrees-of-freedom parameter. Posteriors were sampled with the No-U-Turn sampler; sampling settings, convergence diagnostics and divergence rates are in the Supplementary Material.

### 2.6 Inference

Because the predictors are strongly collinear, a region of practical equivalence applied to an individual coefficient is not a valid decision rule: the probability it computes is conditional on an independence that does not hold. Per-coefficient region-of-practical-equivalence mass (Kruschke, 2018) is therefore reported but explicitly not used as a test. Equivalence statements were made instead on three quantities that are single, well-identified scalars: the coefficient of the global factor; the share of outcome variance carried by the entire local block within the main model; and the difference in Bayesian *R*^2^ between the models with and without that block.

Predictive comparison used leave-one-out cross-validation with Pareto-smoothed importance sampling (Vehtari et al., 2017), with Pareto k diagnostics inspected for every model. Differences in expected log pointwise predictive density were computed pairwise against the global-only model, with standard errors obtained from the pointwise differences rather than from the two marginal standard errors, and a raw difference below four units was treated as small irrespective of its standard error (Sivula et ah, 2025).

### 2.7 Stability analyses

Three resampling analyses distinguished sampler variability from sampling variability. The main model was refitted under three random seeds; refitted on independent halves of the cohort over six random splits; and subjected to stability selection over lasso paths on 200 subsamples of half the cohort, alongside bootstrap sign consistency for every coefficient (Meinshausen and Bühlmann, 2010). A multiverse of 768 specifications crossed the method of intracranial volume adjustment, the age model, inclusion of sex and education, the label set, whether the global factor was removed, hemispheric treatment and winsorisation, with the incremental *R*^2^ of the local block computed by exact leave-one-out rather than in sample (Steegen et ah, 2016; Simonsohn et ah, 2020).

Independently of any outcome, pairs of subregions were classified as exchangeable when entering each alone alongside the global factor changed exact leave-one-out *R*^2^ bv less than 0.002, and equivalence classes were obtained by average-linkage clustering on one minus the absolute correlation.

### 2.8 Planted-effect experiment

The observed data provide no positive signal against which recovery can be judged, so outcomes with a known dependence on the measured volumes were generated. In each repeat, k subregions were drawn at random and assigned coefficients of specified sign and relative magnitude, and the overall scale was solved so that the planted predictors accounted for exactly 2% or *5% of* outcome variance. The identical latent configuration was applied to a ZCA-whitened copy of the same design, which preserves the number of predictors, the sample size and the planted effect size and removes only the correlation among subregions. ZCA rather than principal-component whitening was used because it is the whitening transform closest to the identity, so each whitened column remains the counterpart of the subregion it came from.

Arms were paired within repeat: both designs received the same subsample, the same columns, the same true predictors, the same signs and weights and the same noise vector, with only the scale solved separately so that the target variance share was met in each. Every dataset was analysed with ridge regression, a lasso path and the regularized horseshoe of the main analysis, so that differences between estimators are never confounded with differences between draws.

The main grid comprised four sample sizes (100, 200, 400 and 638), two planted effect sizes and 80 paired repeats per cell, for 1,280 posterior fits with three true predictors of equal magnitude at 36 subregions. A closing round added the parcellation gradient, with 6, 12, 18 and 36 subregions at the observed sample size, and varied the planting scheme over one, three and six true predictors and over equal and decaying magnitude patterns, for a further 1,320 fits.

Recovery curves were summarised by fitting *A*(*n*) = *a — b_n_*^-1/2^, with both parameters bootstrapped by resampling repeats within each sample size. Contrasts between arms were bootstrapped with the pairing preserved, resampling the same repeat indices in both arms. Where a fitted ceiling met the upper estimation bound, a companion fit with the ceiling constrained to one was computed so that the implied sample size could be reported as a range.

### 2.9 Analytic history

Alternative outcomes were specified before the data were analysed, and the framing moved between them according to those pre-specified criteria rather than according to which result was more favourable. One substantive deviation occurred. The identifiabilitv grids were first computed with ridge regression; the estimator comparison described above subsequently showed that ridge, which distributes coefficient mass across correlated predictors instead of selecting among them, recovers the planted set in 24.6% of repeats where the reported model reaches 69.2%. Conclusions resting on those grids were withdrawn and recomputed with the reported model, and the estimator comparison is presented as a result rather than relegated to a sensitivity analysis. A hypothesis that divergent transitions would be more frequent in the collinear design than in the whitened one was tested and not supported, and is reported as such. Dated code and analysis checkpoints for every stage are archived with the data release.

### 2.10 Use of a large language model

A large language model, accessed through a commercial web interface with programmatic handling of its application programming interface, was used during the preparation of this work. The research question and the decision to pursue it were the authors’. The model assisted in the design of the analysis and in the choice of methods, wrote the analysis code, produced drafts of text that the authors subsequently revised, translated the manuscript into English, the authors not being native speakers of that language, and carried out literature searches whose results the authors verified against primary sources. The model had no access to the data and executed no analysis; all computation was run by the authors on the workstation described in Online Resource 1, and the selection and interpretation of results were the authors’ throughout. The authors have verified the whole of the manuscript and the supplementary material against the outputs of that computation and accept responsibility for their content, including the accuracy of every reported value and of every citation.

### 2.11 Software and availability

Analyses were run in Python within a conda environment built for this study (conda 26.3.2), using PyMC 6.0.1 (Abril-Pla et al., 2023), ArviZ 1.3.0 (Martin et al., 2026), NumPy 2.4.6 (Harris et al., 2020), SciPy 1.18.0 (Virtanen et al., 2020) and scikit-learn 1.9.0 (Pedregosa et al., 2011). Full environment and hardware specifications are given in the Supplementary Material. Projection predictive selection, the whitening transform, the planted-effect experiment and the saturation fits were implemented for this study. Reporting follows COBIDAS (Nichols et al., 2017) for the imaging analyses and the Bayesian Analysis Reporting Guidelines (Kruschke, 2021) for the statistical models. Analysis code, configuration files, per-fit outputs and figure-generating scripts are archived and openly available.

## 3 Results

### 3.1 Cohort and design structure

Of the 653 participants with hippocampal subfield segmentations, 638 had complete data on the primary outcome and all covariates and were retained. Participants ranged from 18.6 to 89.0 years of age (mean 55.2, SD 18.4), were evenly divided by sex, and reported a mean of 14.8 years of education (SD 2.8). Estimated intracranial volume averaged 1,600,632 mm^3^ (SD 161,920) and the total modelled hippocampal volume 7,422 mm^3^ (SD 947). CognitiveG had a standard deviation of 1.36 across the cohort.

**Table 1.** Cohort characteristics. Values are mean (standard deviation) and range, for the 638 participants with complete data. Sex is coded 1 for female. CognitiveG and its secondary components are principal components of the cognitive battery and are therefore on an arbitrary scale

| Variable | Mean (SD) | Minimum | Maximum |
| --- | --- | --- | --- |
| Age (years) | 55.16 (18.38) | 18.60 | 89.00 |
| Sex (female / male) | 319 / 319 |  |  |
| Education (years) | 14.83 (2.79) | 9.00 | 18.00 |
| Estimated intracranial volume (mm<sup>3</sup>) | 1,600,632 (161,920) | 1,216,949 | 2,102,245 |
| Total modelled hippocampal volume (mm<sup>3</sup>) | 7,422 (947) | 4,646 | 10,689 |
| CognitiveG | 0.014 (1.362) | -4.442 | 2.949 |
| CognitivePC2 | 0.005 (1.007) | -3.307 | 2.161 |
| CognitivePC3 | 0.002 (0.893) | -3.567 | 2.834 |

The 36 modelled subregions were strongly interdependent (Fig. 1). After adjustment for the nuisance set, the first principal component of the subregion correlation matrix accounted for 42% of its variance, the condition number of the design was 1,727, and the median variance inflation factor was 7.3 with a maximum of 59.3. Bivariate associations with CognitiveG were uniformly weak and almost uniformly positive, ranging from — 0.022 to +0.110, with 32 of 36 correlations positive. Six of the 36 labels, the parasubiculum, fimbria and hippocampus-amygdala transition area in both hemispheres, carry documented reliability limitations (Brown et ah, 2020; Chiappiniello et ah, 2021) and were flagged in advance.

**Fig. 1.**
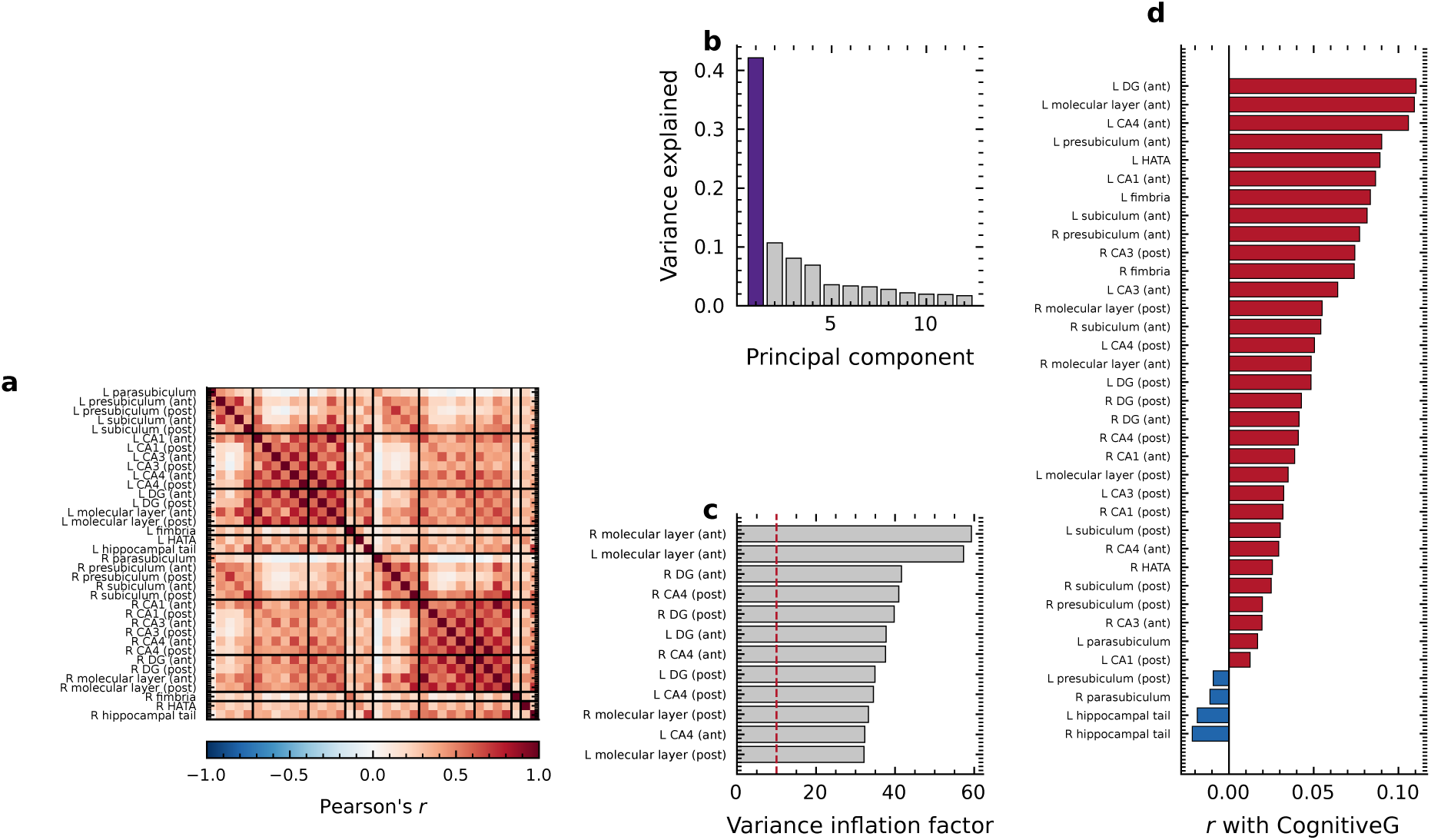
Structure of the hippocampal subregion design (n = 638, p = 36). (a) Pearson correlation between nuisance-adjusted, standardised log volumes, ordered anatomically, with black lines separating the subicular complex, cornu ammonis, dentate and molecular layer, white matter and remaining labels within each hemisphere, (b) Eigenvalue spectrum of that correlation matrix; the first component, in purple, accounts for 42 per cent of the variance, (c) The twelve largest variance inflation factors; the dashed line marks the conventional threshold of ten and the condition number of the design is 1727. (d) Correlation of each subregion with CognitiveG before any multivariate adjustment, sorted; 32 of 36 are positive, so a negative multivariate coefficient would have to be read as suppression rather than as an inverse association.

The nuisance set absorbed a substantial share of variance from both sides of the model, with a median of 0.419 across the subregion volumes (range 0.125 to 0.568) and 53.3% of the variance in CognitiveG, almost all of it attributable to age. The unweighted global factor accounted for a mean of 41.8% of the variance in the adjusted subregion volumes and correlated r = +0.074 with adjusted CognitiveG, explaining 0.6% of its variance on its own. Ten of the 36 subregions lost more than half their bivariate association with the outcome once the global factor was removed (Fig. 2).

**Fig. 2.**
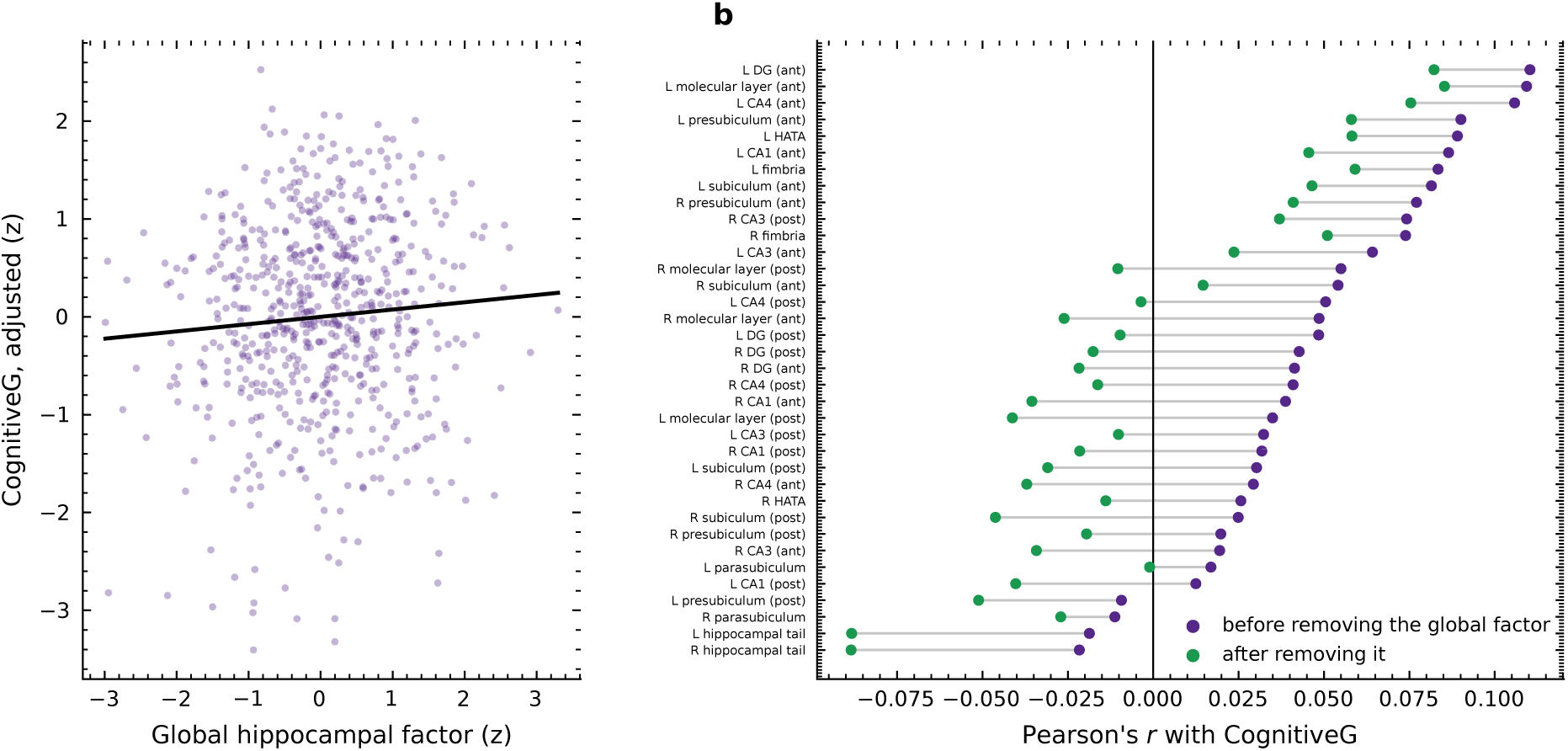
Decomposition of the hippocampal signal into a global component and local contrasts, (a) The unweighted mean of the standardised subregion volumes, the global factor, against adjusted CognitiveG; the correlation is r = +0.074, accounting for 0.6 per cent of outcome variance on its own, with the least-squares line drawn in black, (b) Correlation of each subregion with CognitiveG before removing the global factor, in purple, and after orthogonalising against it, in green, joined by a grey line for each subregion. 10 of 36 subregions lose more than half their association, so most of the apparent regional signal is carried by the common size axis.

The interdependence was strong enough that most pairs of subregions could not be distinguished by the data. Entering each subregion alone alongside the global factor and comparing exact leave-one-out *R*^2^, 58.6% of all pairs differed by less than 0.002 and were therefore exchangeable in the sense that substituting one for the other left out-of-sample fit unchanged (Fig. 3). This quantity is a property of the design and was computed without reference to any outcome.

**Fig. 3.**
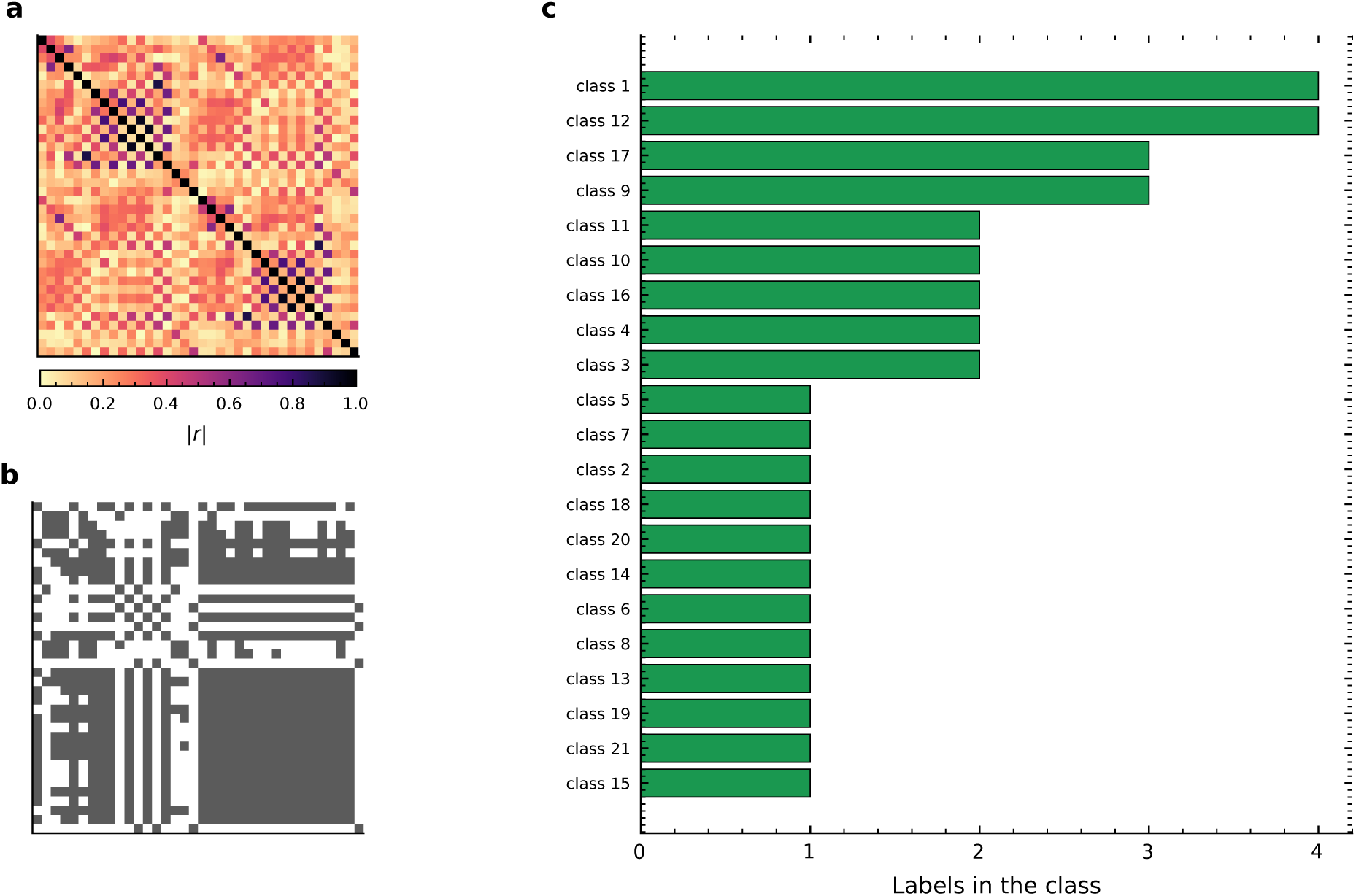
Redundancy of the hippocampal subfield segmentation, computed from the design alone and therefore independent of any outcome, (a) Absolute Pearson correlation between nuisance-adjusted subregion volumes, in the same anatomical order as Figure 1, where the labels are given, (b) Pairs of subregions that are exchangeable, shown in black, defined as differing by less than 0.002 in exact leave-one-out R^2^ when each is entered alone alongside the global factor; 59 per cent of all pairs qualify, so for most pairs the data cannot prefer one member over the other, (c) Sizes of the 21 equivalence classes obtained by average-linkage clustering at |r| *≥* 0.6, the largest containing 4 labels. A result should therefore be reported at the level of a class rather than of an individual label.

### 3.2 No subregion was identifiable in the observed data

Under the regularized horseshoe, no local coefficient was distinguishable from zero. None of the 36 equal-tailed intervals excluded zero, the posterior mean of the effective number of non-zero coefficients was 1.89 against a prior guess of four, and no pair of coefficients showed the posterior anticorrelation that would indicate the model trading one label against another (Fig. 4). The unshrunk global coefficient was +0.067 with an 89% equal-tailed interval of [0.004,0.130], so that its interval excluded zero while a third of its posterior mass lay inside the region of practical equivalence.

**Fig. 4.**
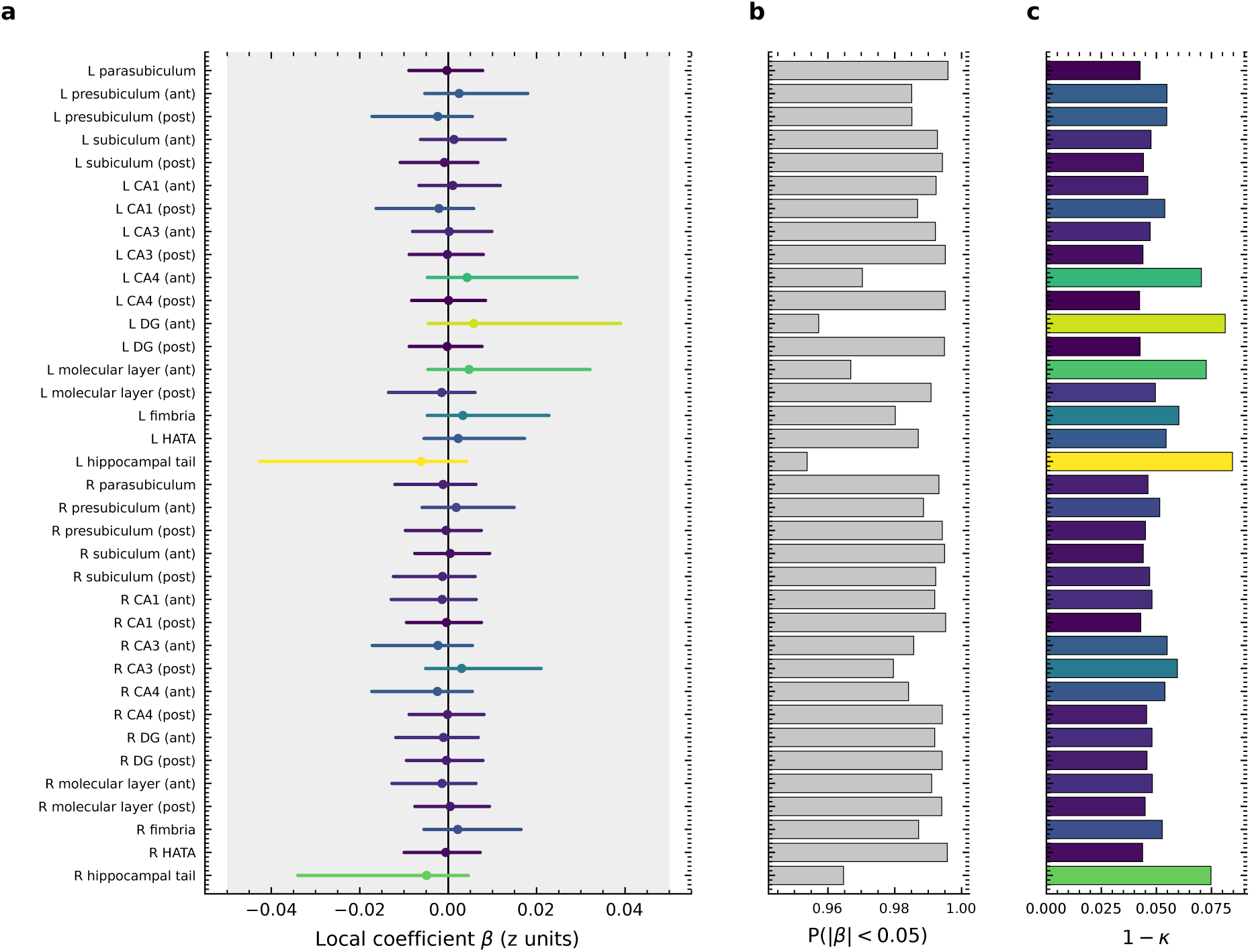
Posterior local coefficients under the regularized horseshoe, with the global factor entered unpenalised at +0.067 [+0.004, +0.130]. (a) Posterior mean and 89 per cent equal-tailed interval for each subregional contrast; colour encodes the posterior shrinkage factor *k,* filled markers would indicate an interval excluding zero and red rings a coefficient whose sign opposes its own marginal correlation, (b) Posterior mass inside the region of practical equivalence. This panel is shown for completeness and is deliberately not used as a test: with predictors this collinear. a region of practical equivalence applied to a univariate marginal is conditional on an independence that does not hold, (c) The fraction of each coefficient that escaped shrinkage, 1 — *κ*, on an axis clipped just above its largest value; the whole range spans 0.042 to 0.085, so no coefficient escapes to any meaningful degree. Subregions with intervals excluding zero: none.

The model ladder separated poorly (Fig. 5). Posterior Bayesian *R*^2^ was 0.005 for whole hippocampal volume alone, 0.007 for the global factor alone, 0.012 for the global factor with the local block and 0.007 for a flat horseshoe over all subregions. The share of outcome variance carried by the local block within the main model was 0.005, with an 83.5% posterior probability of falling below 0.01, and the difference in Bayesian *R*^2^ between the model with and without the local block was +0.005 with an 89% interval of [—0.012, 0.024]. No pairwise difference in expected log pointwise predictive density against the global-only model exceeded four units in magnitude, the threshold below which a difference is treated as small irrespective of its standard error. The posterior probability that the difference in Bayesian *R*^2^ fell inside the region of practical equivalence was 0.64, which is insufficient to assert equivalence; the result is non-detectability rather than demonstrated absence.

**Fig. 5.**
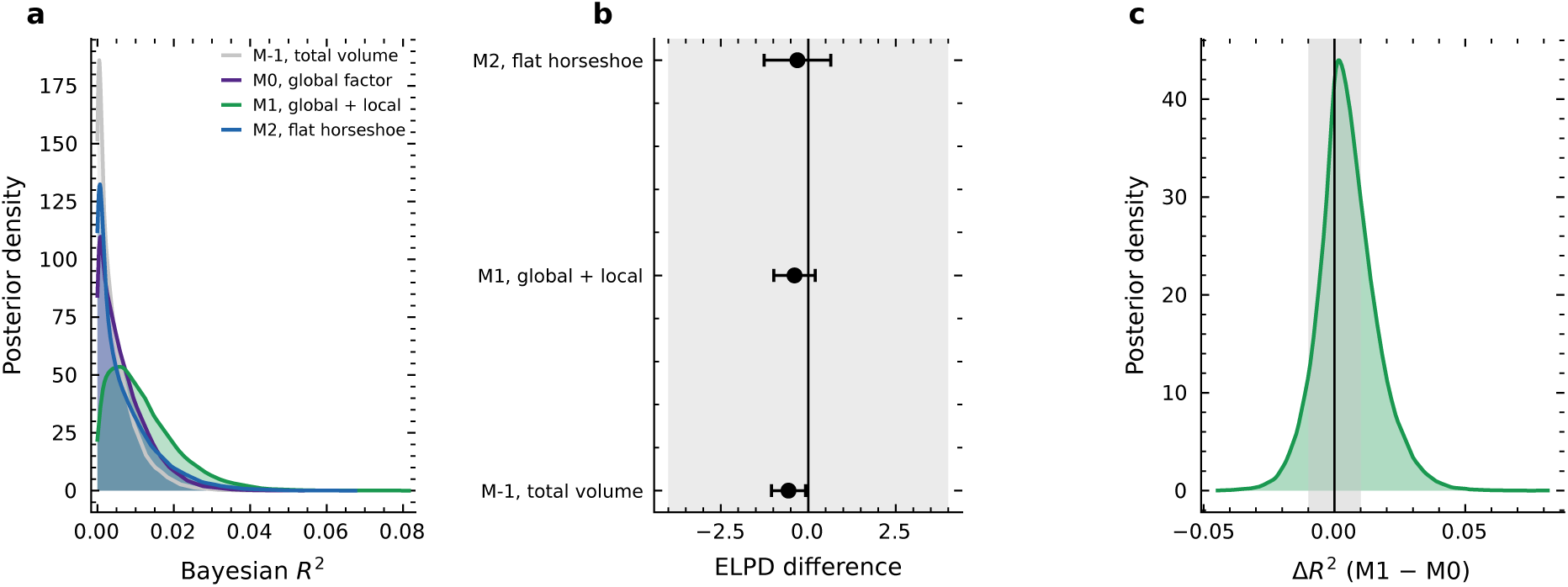
Whether subregional topography adds anything beyond global size. (a) Posterior Bayesian R2 for each model in the ladder, with posterior means of M-1, total volume 0.005, M0, global factor 0.007, M1, global + local 0.012, M2, flat horseshoe 0.007. (b) Difference in expected log pointwise predictive density from leave-one-out cross-validation against the global-factor model, with standard errors computed from the pointwise differences rather than from the two marginal standard errors; the shaded band spans differences of less than 4 units, which are treated as small irrespective of their standard error. (c) Posterior difference in Bayesian *R*^2^ between the models with and without the local block, with the region of practical equivalence shaded; 0.640 of the posterior mass falls inside it. Unlike a region of practical equivalence applied to an individual collinear coeffcient, this one is applied to a single well-identified scalar and is a valid equivalence statement.

Two resampling analyses separated sampler noise from sampling noise (Fig. 6). Refitting the same model on the same data under three random seeds returned the same answer each time: posterior means fall on the identity line, and the mean pairwise Jaccard overlap among the top six coefficients by absolute posterior mean is 1.00. Refitting on independent halves of the same cohort did not. Across six random splits the mean correlation between the two halves’ coefficient vectors was +0.12, with a mean top-k overlap of 0.08, and the per-split detail is given in Fig. 7. Across 768 analytic specifications the global coefficient remained stable in sign and magnitude while twelve different subregions took the position of largest local coefficient, and 576 of those specifications produced a negative out-of-sample gain from the local block.

**Fig. 6.**
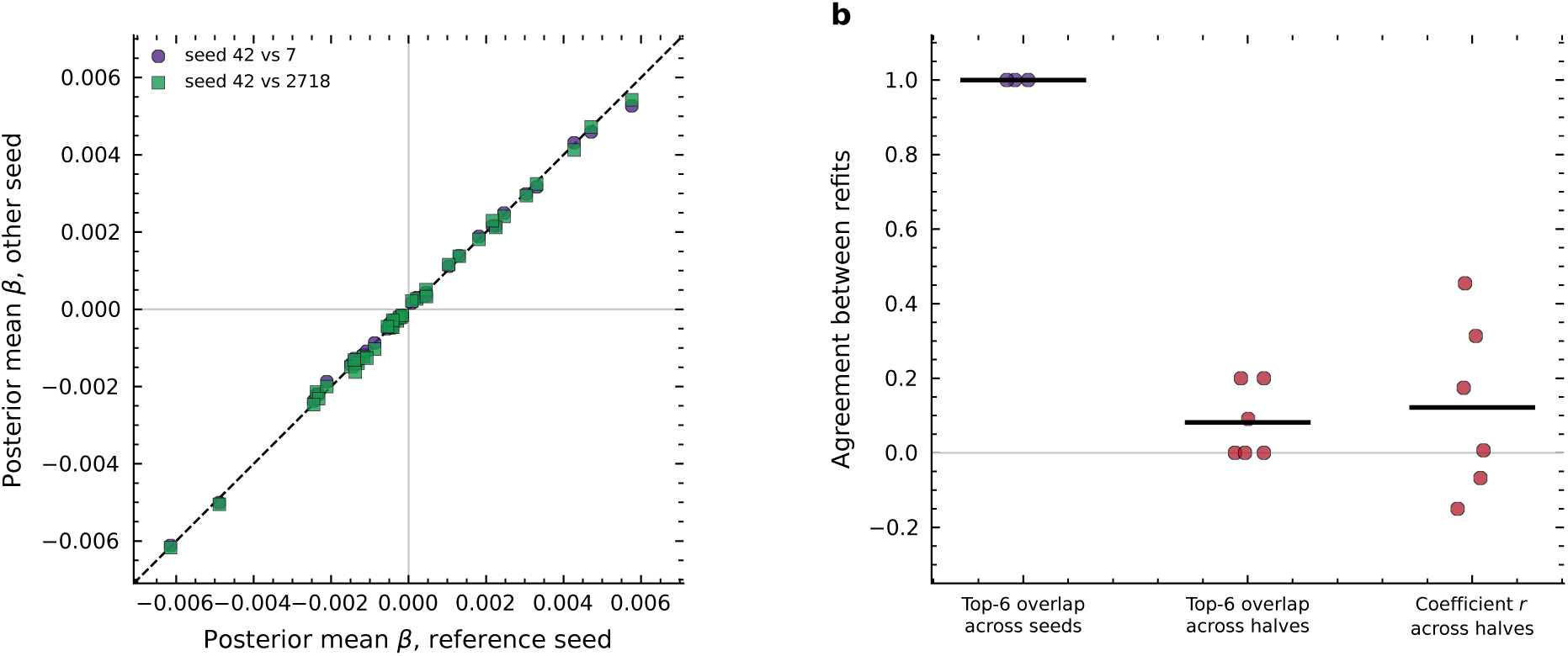
Sampler variability against sampling variability, (a) Posterior mean coefficients from refits of the same model on the same data, differing only in the random seed, plotted against the reference seed: the dashed line is the identity. Points fall on it. so the estimator returns the same answer every time it is asked, (b) Agreement between refits, with each point one comparison and the horizontal bar its mean: overlap of the top six selections across seeds (mean 1.00). the same overlap across independent halves of the cohort (mean 0.08). and the correlation between the two halves’ coefficient vectors (mean +0.12). Refits that share the data agree perfectly: refits that share only the cohort do not. The instability is in the estimand. not in the estimation.

**Fig. 7.**
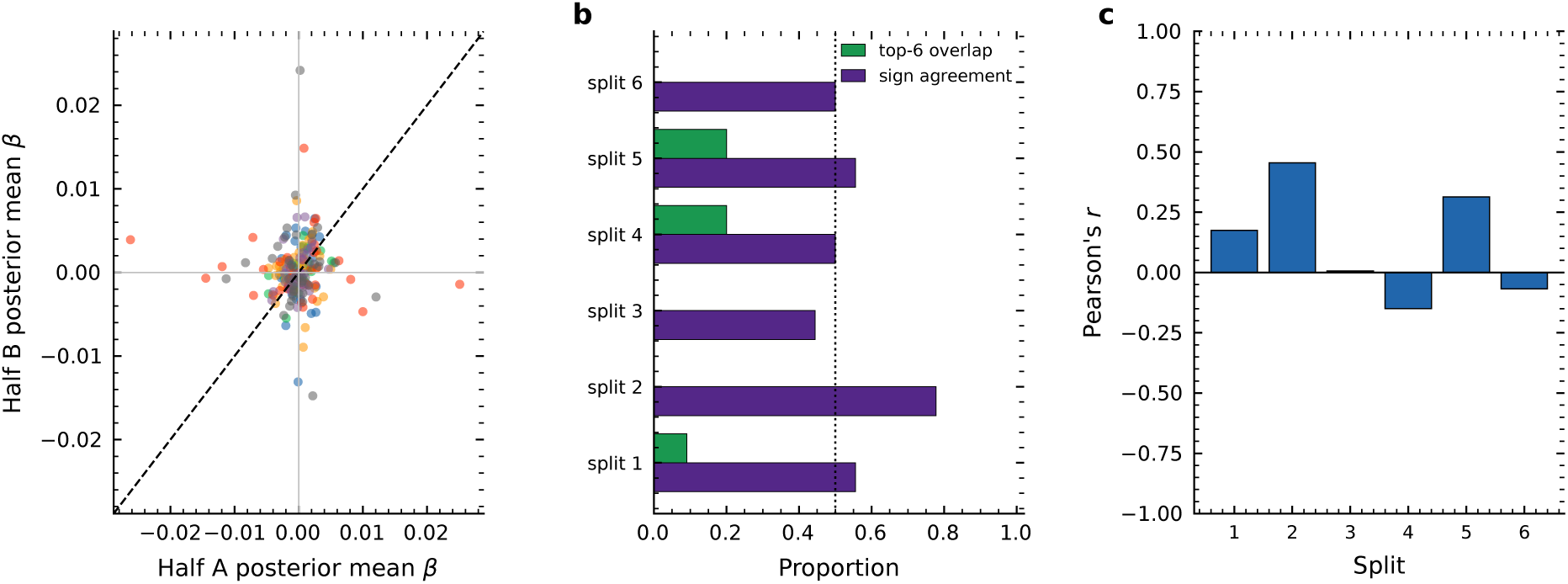
Reproducibility of the subregional pattern between independent halves of the same cohort, over 6 random splits, with the full model refitted on each half, (a) Posterior mean coefficients from one half against the other, one colour per split: agreement would place points on the dashed identity line, (b) For each split, the overlap of the top-6 selections and the proportion of coefficients whose sign matches across halves: the dotted line marks the value expected if signs were assigned at random, (c) Pearson’s correlation coefficient between the two halves’ full coefficient vectors, one bar per split, with a mean of +0.122. Two halves of one cohort share scanner, protocol and population, so this is an upper bound on what an independent replication could achieve.

Five further analyses specified in the Methods are reported in full in the supplementary material, and none alters the conclusion of this section. Refitting after removing the six labels with documented reliability limitations leaves 30 predictors and no interval excluding zero; replacing the local contrasts with centred log-ratio coordinates gives 36 coordinates and, again, none excluding zero. Stability selection over 200 subsamples reaches a highest selection frequency of 0.515, below the 0.60 threshold, and the median sign consistency across bootstrap resamples is 0.772. Projection predictive selection gains 1.8 units of expected log pointwise predictive density in sample and 0.08 under cross-validation. A multivariate model over the three cognitive components places the difference between the general factor and each secondary component at +0.053 and +0.050, with intervals including zero, so the association is not demonstrably specific to general cognitive ability.

### 3.3 Recovery of a known effect under the real and orthogonalised designs

Because the observed data provided no positive signal against which recovery could be judged, we generated outcomes with a known dependence on the measured subregion volumes. In each repeat, three of the 36 subregions were drawn at random and assigned coefficients of equal magnitude and random sign, scaled so that the planted predictors accounted for exactly 2% or 5% of outcome variance. The same procedure was applied to a ZCA-whitened copy of the design, which preserved the number of predictors, the sample size and the planted effect size while removing the correlation among subregions. Arms were paired: within a repeat, both designs received the same subsample, the same columns, the same true predictors, the same signs and the same noise vector, and only the coefficient scaling was solved separately so that the target variance share was met in each. Every dataset was analysed with the regularized horseshoe used in the primary analysis, under identical priors. The grid comprised four sample sizes (n = 100, 200, 400, 638) and 80 paired repeats per cell, for 1,280 posterior fits.

Recovery of the complete planted set increased monotonically with sample size in both arms and at both effect sizes (Fig. 8; full series in Supplementary Table S3). At a planted effect of 5%, the real design recovered all three planted subregions in 20.8% of repeats at *n* = 100 and 69.2% at *n* = 638, against 30.0% and 85.0% for the whitened design. At a planted effect of 2% the corresponding values were 15.4% and 38.3% for the real design and 15.4% and 53.3% for the whitened one. Fitting agreement(n) = a — bn^-1/2^ to each curve, with both parameters bootstrapped over repeats, gave a real-design ceiling of 0.944 (89% interval 0.878 to 1.009) at a 5% effect and 0.544 (0.483 to 0.603) at a 2% effect; the whitened ceilings were 1.05 at the upper estimation bound and 0.738 (0.681 to 0.800) respectively (Supplementary Table S4).

**Fig. 8.**
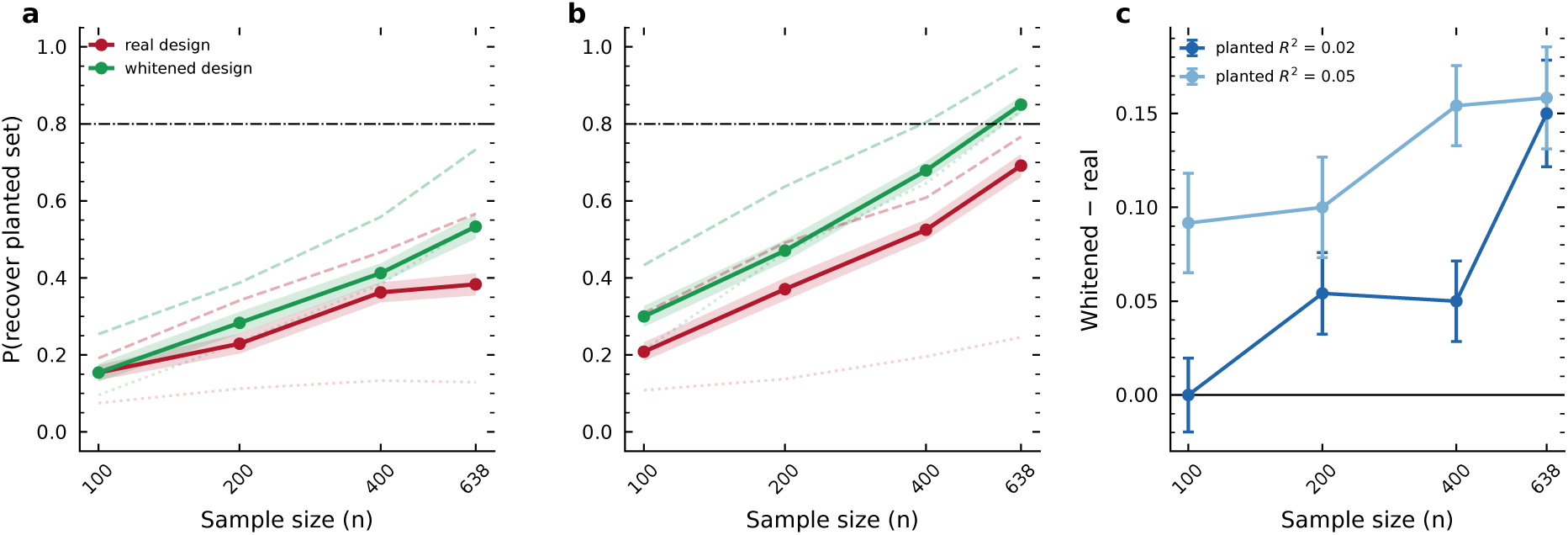
Recovery of a planted effect by the regularized horseshoe reported in the article, in the measured design and in a ZCA-whitened copy of it that preserves sample size, dimensionality and effect size while removing only the correlation between subregions. Within each repeat the two arms receive the same subsample, columns. 3 true predictors, signs and noise vector, so they differ only in the covariance of the design, (a) planted effect of 0.02 of outcome variance, (b) planted effect of 0.05 of outcome variance. The horseshoe is drawn solid with markers and the lasso and ridge faintly behind, dashed and dotted, because the estimator was found to change recovery by a factor of three: bands are standard errors over 80 paired repeats and the dash-dotted line is the recovery criterion of 0.8. (c) The paired whitened minus real difference with its own standard error, which is the quantity separating a limit imposed by the geometry from one imposed by sample size.

### 3.4 Collinearity imposes a sample-size cost rather than a ceiling

Neither arm was bounded away from adequate recovery at the larger planted effect: both reached the 0.80 criterion, and the difference between them was expressed in the sample size required to do so. Because the whitened asymptote met the upper estimation bound, the requirement implied by the unconstrained fit is a lower bound for that arm, and a companion fit with the ceiling held at one was computed to obtain the other end of the range. Taken together the two fits placed the requirement at 1,752 to 2,803 participants for the real design and 893 to 1,151 for the whitened design at a 5% planted effect, so that collinearity among subregions multiplied the required sample size by a factor between 1.5 and 3.1 without rendering the target unattainable. The same companion fit was not admissible at the smaller planted effect, where both fitted ceilings excluded one (0.544, 89% interval 0.483 to 0.603 for the real design; 0.738, 0.681 to 0.800 for the whitened design) and a requirement computed under an assumed ceiling of one would contradict the data it was fitted to.

The paired construction allowed this difference to be estimated directly rather than by comparing independently fitted curves. The whitcncd-minus-rcal difference in recovery excluded zero in seven of eight cells (Fig. 8c, Supplementary Table S3). At a 5% planted effect it was 0.092 ± 0.027 at *n* = 100 and 0.158 ± 0.027 at *n* = 638; at a 2% planted effect it was 0.000 ± 0.020 and 0.150 ± 0.028. The single cell consistent with zero was the smallest sample at the smallest effect, where neither arm recovered the planted set in more than 16% of repeats.

### 3.5 The cost of collinearity increases with sample size

The paired difference was not constant across the sampling range, and its behaviour depended on effect size. In absolute terms the separation between arms widened with *n* in both conditions, from 0.092 to 0.158 at a 5% planted effect and from 0.000 to 0.150 at a 2% effect. Expressed relative to the recovery achieved by the whitened design, the two conditions diverged: the shortfall of the real design fell from 31% to 19% at the larger effect while rising from 0% to 28% at the smaller one.

The saturation fits located this divergence in different parameters. At a 5% planted effect the arms approached their asymptotes at indistinguishable rates, with a paired whitened-minus-real difference in b of — 0.13 (89% interval —1.01 to 0.78), while their ceilings differed by 0.106 (0.043 to 0.171), a difference that reads as a lower bound because the whitened asymptote met the upper estimation bound. At a 2% planted effect both parameters differed: b by 2.01 (1.20 to 2.79) and the ceiling by 0.194 (0.134 to 0.252). At the larger planted effect the two designs therefore converted additional observations into recovery at the same rate but toward different limits, whereas at the smaller effect the real design was slower to improve as well as bounded lower.

### 3.6 At realistic effect sizes, orthogonality is not sufficient either

Both designs failed to reach the recovery criterion at the smaller planted effect. At a 2% planted effect the real design recovered the complete planted set in 38.3% of repeats at *n* = 638 and the whitened design in 53.3%, with fitted ceilings of 0.544 (89% interval 0.483 to 0.603) and 0.738 (0.681 to 0.800) respectively. Neither interval included the 0.80 criterion, and under the fitted saturation model neither configuration reached it at any sample size. Removing the correlation among subregions therefore raised recovery substantially without making the target attainable.

Reducing the number of units did not rescue it either. At the same effect size and sample size, recovery in the real design reached only 0.750 ± 0.028 at the coarsest parcellation tested, six subregions, and the whitened design 0.783 ± 0.025, so that neither design met the criterion even when the number of candidate units was reduced sixfold and the correlation among them removed entirely (Fig. 9a). At this effect size the limitation is therefore not attributable to the geometry of the parcellation, to its granularity, or to either in combination, but to the magnitude of the effect relative to the sample size available.

**Fig. 9.**
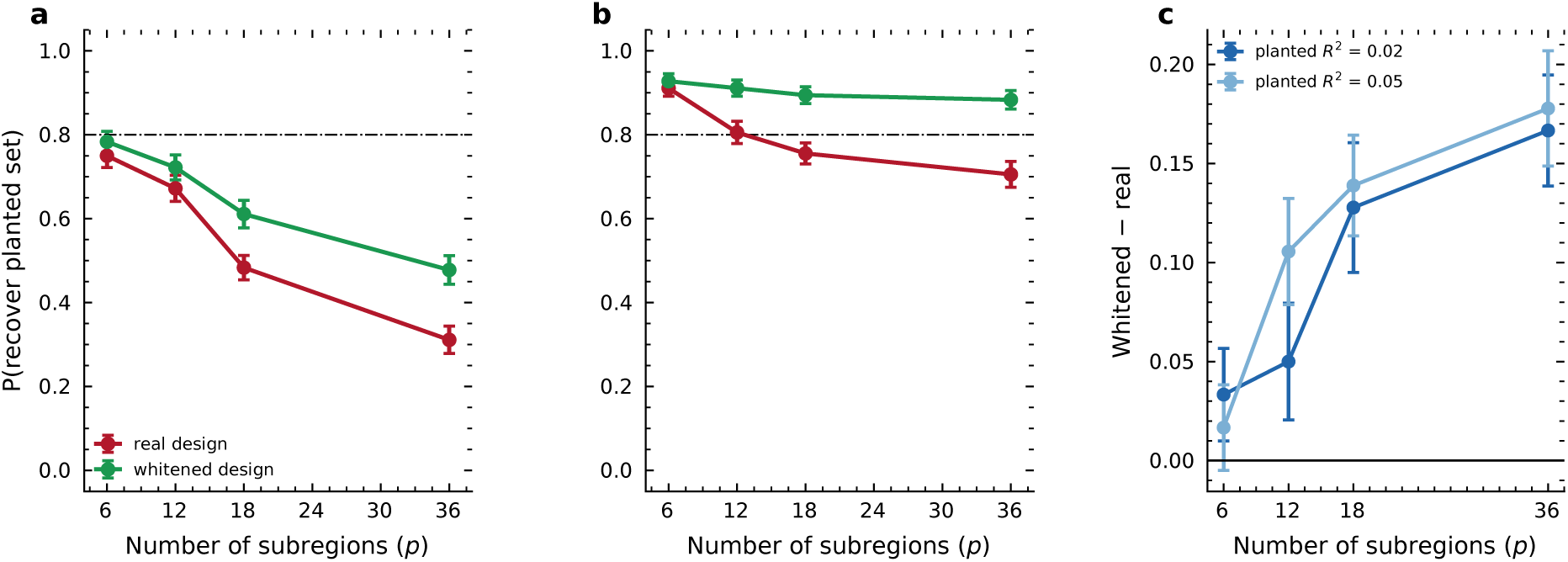
Recovery of a planted effect against parcellation granularity, measured with the regularized horseshoe at the observed sample size of n = 638. In each repeat p subregions are drawn at random from the 36 available, 3 of them carry the planted effect, and the identical configuration is applied to the measured design and to its whitened counterpart, (a) planted effect of 0.02 of outcome variance, (b) planted effect of 0.05 of outcome variance. Error bars are standard errors over 60 paired repeats and the dash-dotted line is the recovery criterion of 0.8. In panels a and b the vertical axis is the probability that all 3 planted subregions appear among the 3 largest estimated coefficients, and the horizontal axis carries ticks at regular intervals although recovery was measured at 6, 12, 18 and 36 subregions only, (c) The whitened minus real difference in that probability, paired within repeat, at each granularity, which isolates how much of the loss is attributable to correlation among the units rather than to their number; at the coarsest parcellations it is consistent with zero, so there the limit is dimensionality and not collinearity.

### 3.7 Recovery depends strongly on the estimator

The three estimators applied to identical data diverged widely (Fig. 10; Supplementary Table S1). At *n* = 638 in the real design with a *5%* planted effect, the complete planted set was recovered in 24.6% of repeats by ridge, 69.2% by the regularized horseshoe and *76.7%* by the lasso path; at a 2%; planted effect the corresponding values were 12.9%;, 38.3% and 56.7%;. Paired differences within repeat placed the horseshoe 0.446 ± 0.038 above ridge and 0.075 ± 0.022 below the lasso at the larger effect, and 0.254 ± 0.033 above ridge and 0.183 ± 0.028 below the lasso at the smaller one.

**Fig. 10.**
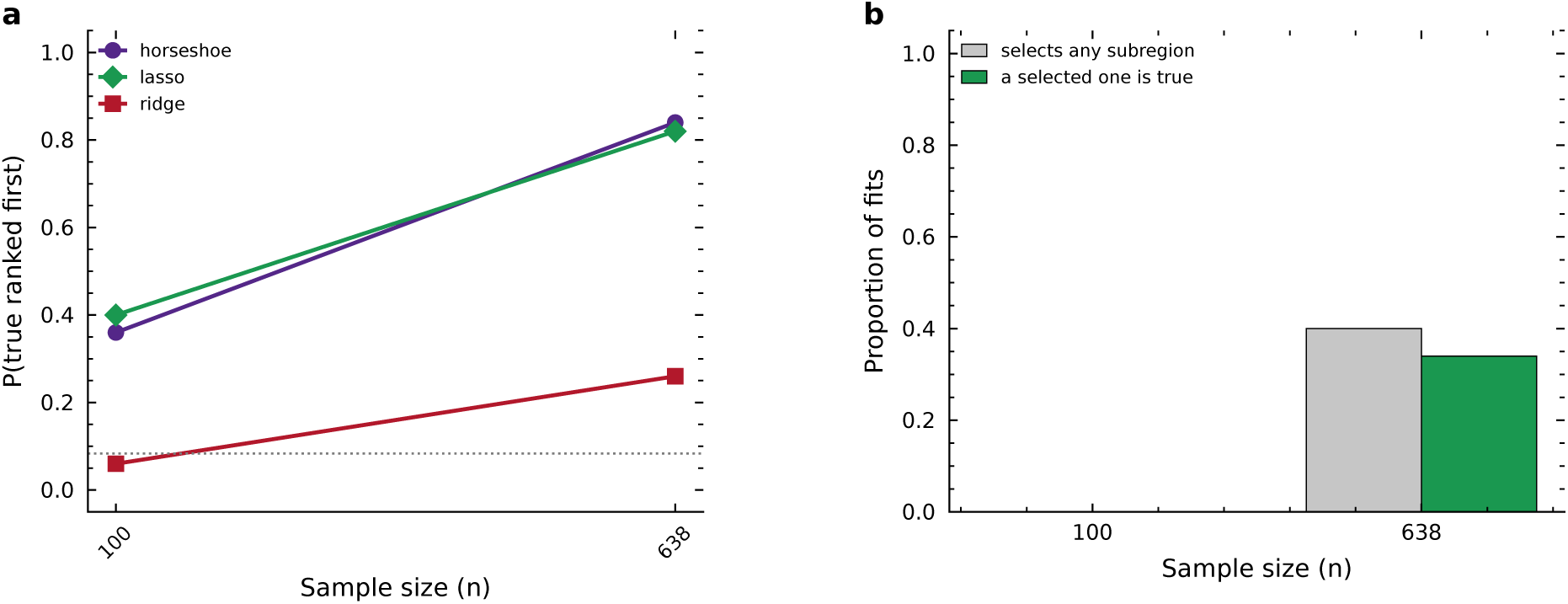
The model reported in the article placed among the estimators used to build the identifiability curves. Every repeat plants the same effect in the same subsample and the same columns and hands it to all three estimators, so the comparison is paired. (a) Probability that a genuinely non-zero subregion is ranked first, at p = 36 and a planted effect of 0.05 of outcome variance, for ridge, for the first entry of a lasso path and for the regularized horseshoe; the dotted line is chance, 3/36. (b) For the horseshoe alone, how often any coeffcient interval excludes zero and how often at least one such coeffcient is genuinely non-zero. The gap between the two bars is the rate at which the reported analysis would name a subregion that carries no planted effect.

The ordering was not preserved across criteria. When the criterion was whether the highest-ranked subregion was genuinely non-zero, the horseshoe was highest at both effect sizes (85.0%; and 56.2%;, against 76.2%; and 51.2%; for the lasso and 28.8%; and 13.8%; for ridge). Ridge, which distributes a coefficient across correlated predictors rather than selecting among them, was the lowest performer under every criterion and in every condition.

### 3.8 The reported model rarely selects, and selects correctly when it does

Selection was defined as a coefficient whose 89% posterior interval excluded zero. In the real design the horseshoe selected no subregion at all in the majority of fits: at a *2%* planted effect it selected nothing in 100% of repeats at *n* = 100 200 and 638 and in 96.2% at *n* = 400. At a 5% planted effect selection became more frequent with sample size, from 0% at *n* = 100 to 5.0% at *n* = 200, 15.0% at *n* = 400 and 41.2% at *n* = 638, with a mean of 0.50 subregions selected per fit at the largest sample. When the model did select, the selection contained a genuinely non-zero subregion in 38.8% of all repeats at *n* = 638, that is in 94.2% of the repeats in which any selection occurred. The corresponding figures for the whitened design were 66.2%; selecting and 66.2%; correct.

### 3.9 Recovery declines with parcellation granularity

Because the parcellation gradient carries the study’s only design recommendation and had been estimated with a lasso path, it was re-measured with the regularized horseshoe at the observed sample size of *n* = 638, drawing *p* subregions at random from the 36 available and planting three true effects among them (Fig. 9, Supplementary Table S5). Recovery of the complete planted set fell monotonically with granularity in every condition. In the real design at a 5% planted effect it declined from 0.911 ± 0.019 at *p* = 6 to 0.806 ± 0.027 at *p* = 12 0.756 ± 0.025 at *p* = 18 and 0.706 ± 0.031 at *p* = 36; at a 2% planted effect the decline was steeper, from 0.750 ± 0.028 to 0.672 ± 0.031 0.483 ± 0.029 and 0.311 ± 0.033. The whitened design declined more gently over the same range, from 0.928 to 0.883 at the larger effect and from 0.783 to 0.478 at the smaller one.

The paired difference between arms grew with granularity and was resolved only beyond the coarsest parcellations (Supplementary Table S6). At a 5% planted effect the whitened-minus-real difference was 0.017 ± 0.022 at *p* = 6 0.106 ± 0.027 at *p* = 12 0.139 ± 0.025 at *p* = 18 and 0.178 ± 0.029 at *p* = 36; at a 2% planted effect it was 0.033 ± 0.023, 0.050 ± 0.029, 0.128 ± 0.033 and 0.167 ± 0.028. Of the eight comparisons, the two coarsest at each effect size were consistent with zero at a 2% planted effect and the coarsest alone at a 5% effect, while every comparison at *p* ≥ 18 excluded zero. Against the 0.80 criterion, the real design met it at *p* = 6 and *p* = 12 at a *5%* planted effect and at no granularity at a *2%* planted effect.

### 3.10 Recovery depends on the number and magnitude pattern of the planted effects

The planted scheme used three true predictors with coefficients of equal magnitude, and both choices were varied at *n* = 638, *p* = 36 and a *5%* planted effect (Fig. 11, Supplementary Table S7). Recovery in the real design was 0.983 ± 0.017 with a single true predictor, 0.692 ± 0.030 with three of equal magnitude, 0.511 ± 0.027 with three whose coefficients stood in the ratio 1,1/2,1/3, and 0.300 ± 0.015 with six of equal magnitude. The whitened design followed the same ordering at 0.983, 0.850, 0.628 and 0.422. The largest deviation from the configuration used for the reported result was 0.392, obtained with six true predictors.

**Fig. 11.**
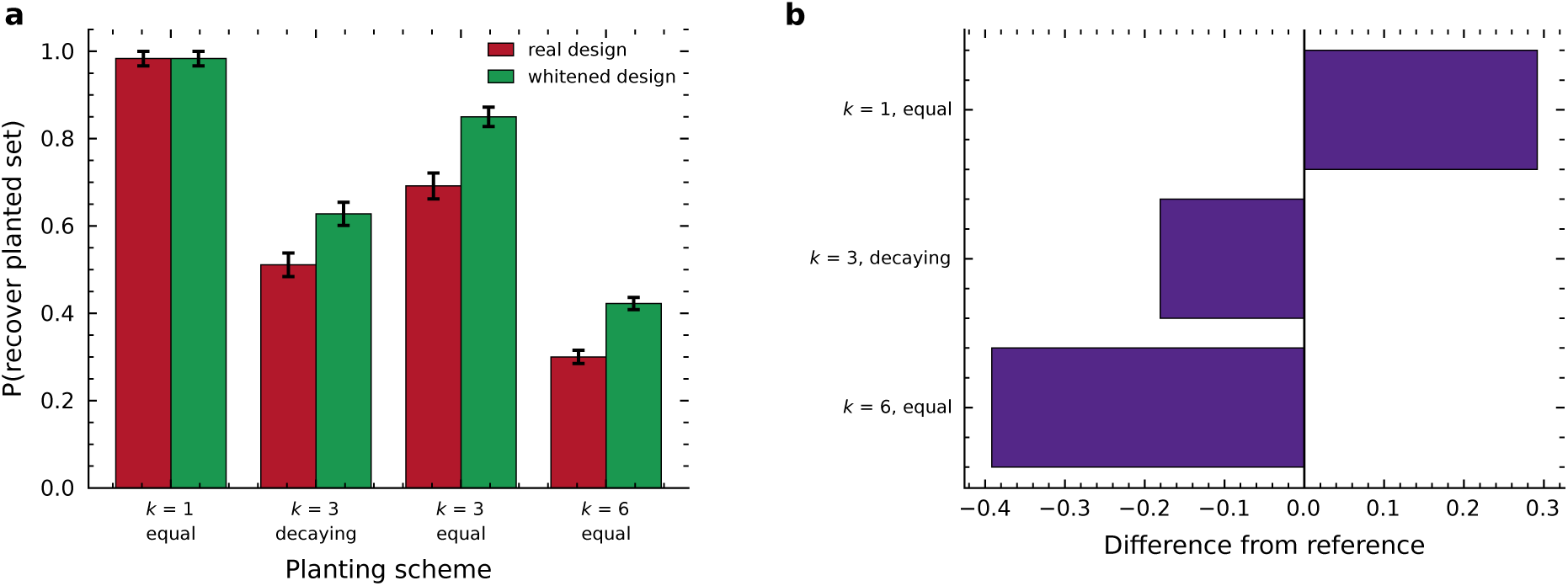
Sensitivity of recovery to the way the effect was planted, at n 638, p 36 and a planted effect of 0.05 of outcome variance. The reported experiment used three true predictors of equal magnitude: both the number and the magnitude pattern were choices rather than findings, so both are varied here, (a) Probability of recovering the complete planted set under one. three and six true predictors of equal magnitude and under three whose coefficients stand in the ratio 1. 1/2, 1/3, for the measured design and its whitened counterpart, with standard errors, (b) Difference of each alternative scheme from the configuration the reported number came from: the largest deviation is 0.392, which exceeds the paired cost of collinearity at the same cell. The reported figure is therefore one point on a calibration surface and not an estimate of what a real study would achieve.

Recovery therefore depended on the planting scheme by considerably more than it depended on the presence of collinearitv, whose paired contribution at this cell was 0.158 ± 0.027. The whitened-minus-real difference was itself smallest where recovery was near ceiling with one true predictor (0.000), and comparable to the main configuration under the decaying pattern (0.117) and under six true predictors (0.122). Both blocks were sampled without failure: no fit of the 1,320 failed to complete, and the median divergence rate was 0.0202 in both arms (Fig. 12).

**Fig. 12.**
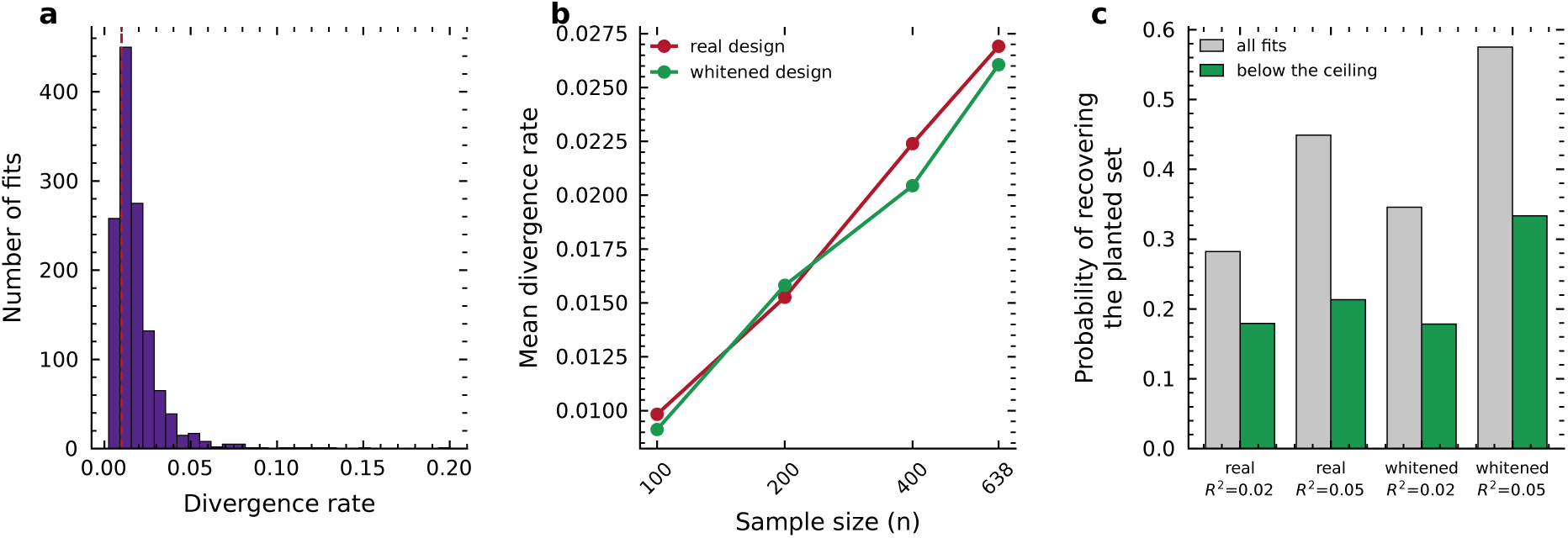
Sampling health of the horseshoe fits behind the planted-effect experiment, (a) Distribution of the proportion of divergent transitions per fit, with the dashed line at the 1% ceiling used below; the median rate is 0.0146. Divergences are expected rather than surprising here, since the horseshoe posterior is multimodal under correlated predictors, (b) Mean divergence rate by design and sample size. The two arms are indistinguishable and the rate scales with sample size, so the divergences do not carry the signature of collinearity we had predicted, (c) Recovery computed on all fits against recovery computed only on fits below the ceiling; agreement between the pairs of bars means the reported recovery does not depend on the divergent fits.

## 4 Discussion

### 4.1 A literature that does not converge

The inconsistency of the hippocampal subfield literature is long-standing and predates subfield segmentation itself. Van Petten’s meta-analysis of 33 studies concluded that a volume--memory relationship probably does not exist among young adults and may emerge only in samples with a higher proportion of older participants (Van Petten, 2004). A later meta-analysis in typically developing children and adolescents reported an association in the opposite direction, with larger hippocampal volume accompanying better memory (Botdorf et al., 2022). Two quantitative reviews of the same construct disagreeing on sign is not a result that any single study’s design explains.

Our observed data reproduce the pattern rather than resolving it. No coefficient interval excluded zero across 36 subregions, the local block carried 0.5% of outcome variance, and no model in the ladder improved out-of-sample predictive fit over a single global factor by more than four units of expected log density. Read alone, that null is uninformative: it is equally consistent with an absent effect and with a present effect the design cannot locate. The two possibilities have opposite implications for how the field should proceed, and distinguishing them requires knowing what the design would do if an effect were there.

### 4.2 Collinearity sets a price, not a wall

Planting an effect of known location answered that question. At a planted effect accounting for 5% of outcome variance, the correlation structure of the subregion design did not prevent recovery of the planted set. It raised the sample size required to achieve it, from between 893 and 1,151 participants in an orthogonalised copy of the same design to between 1,752 and 2,803 in the real one, a multiplier between 1.5 and 3.1.

That multiplier is not a fixed offset. The paired difference between arms widened as sampling grew, from 0.092 at *n* = 100 to 0.158 at *n* = 638 at a 5% planted effect, and the saturation fits located the divergence in different parameters depending on effect size. At 5% the arms approached their limits at indistinguishable rates and differed only in where they stopped. At 2% the real design was also slower to improve, with a paired difference in the approach rate of 2.01 (89% interval 1.20 to 2.79). Each additional participant therefore bought less recovery in the correlated design than in the orthogonal one, and the shortfall compounded rather than closing.

At the effect sizes this literature reports, none of this was sufficient. With a 2% planted effect the real design recovered the complete set in 38.3% of repeats at the full sample and the whitened design in 53.3%, with fitted ceilings of 0.544 and 0.738. Neither reached the 0.80 criterion at any sample size. Coarsening the parcellation did not rescue it either: at six subregions rather than 36, recovery reached only 0.750 in the real design and 0.783 in the whitened one. Here the binding constraint is the size of the effect relative to the cohort, and collinearity adds to a bill the study could not pay in any case.

### 4.3 Detection and localisation do not scale together

The argument that brain-wide association studies require thousands of participants rests on the observation that the largest replicable associations reached r = 0.14 univariately and r = 0.34 multivariatelv (Marek et ah, 2022). That conclusion has been contested. Multivariate models have been shown to replicate with substantially smaller samples in favourable conditions (Spisak et ah, 2023), prompting a reply from the original authors (Tervo-Clemmens et ah, 2023), and a parallel line of argument holds that design choices can establish brain-behaviour relationships without large enrolment (Rosenberg and Finn, 2022; Gratton et ah, 2022).

Both positions concern whether an association reproduces. Neither addresses whether its anatomical location does, and in our experiment the two came apart. At the full sample and a 5% planted effect, a sparse selector named a genuinely non-zero subregion first in 76.2% of repeats while the complete planted set was recovered in 69.2%; at 2% the figures were 51.2% and 38.3%. A study can be adequately powered to report that hippocampal volume relates to cognition and underpowered to say which part of the hippocampus carries the relationship. A power calculation built on detection will not reveal this, because the quantity it certifies is not the quantity the paper’s conclusion depends on.

### 4.4 A statistical account, and why the estimator mattered

The behaviour has a known form in high-dimensional statistics. Support recovery bv l_1_ methods requires an ir represent able condition on the design, which fails when relevant and irrelevant predictors are strongly correlated (Zhao and Yu, 2006), with neighbourhood-stability conditions playing the same role (Meinshausen and Bühlmann, 2006). Recovery thresholds have been derived in which the required sample size scales as a design-dependent constant multiplying klog(*p* — *k*), the constant being a function of the predictor covariance (Wainwright, 2009). Our paired comparison is that dependence made empirical. Holding *k, p, n* and effect size fixed and changing only the covariance moved the requirement by a factor of 1.5 to 3.1. We did not compute the covariance-specific constants themselves, so the correspondence is qualitative.

The same account explains a result that could otherwise be mistaken for a property of the data. Ridge regression, which spreads coefficient mass across correlated predictors instead of choosing among them, recovered the planted set in 24.6% of repeats where the regularized horseshoe reached 69.2% and the lasso path 76.7%. Any of the three could have been chosen in advance without comment, and the choice would have moved the headline by a factor of three. Adaptive weighting schemes that relax the condition of Zhao and Yu (2006) are known to improve support recovery (Zou, 2006), and the ordering we observed follows that theory.

Posterior multimodality under correlated predictors, documented for the horseshoe (Piironen and Vehtari, 2017), also accounts for the sharpest contrast in the observed data. Coefficient solutions agreed perfectly across sampler seeds, with a top-k Jaccard of 1.00, and correlated at r = 0.12 between independent halves of the same cohort. The estimator is stable. The estimand is not.

### 4.5 The genetics convention as external validation

Genetic studies of these structures adopted a convention the cognitive literature never standardised. In the largest such analysis, association testing was run on each subfield while co-varying for whole hippocampal volume, stated explicitly as a means of identifying signal specific to one or some of the subfields (van der Meer et ah, 2020). The convention concedes, in practice, that subfield volumes are not separable from the global factor unless the global factor is removed by hand. Our global and local decomposition performs the same operation on a cognitive outcome, and the planted-effect experiment prices it. The apparent contrast between a genetics literature that reports subfield-specific signal and a cognitive literature that does not owes something to one field having settled the adjustment and the other having left it to the analyst, on top of an order-of-magnitude difference in sample size.

### 4.6 What follows for practice

Granularity is a decision with a measurable cost. Recovery in the real design fell from 0.911 at six units to 0.706 at 36 with a 5% planted effect, and from 0.750 to 0.311 at 2%. The paired cost of collinearity was resolved only from 18 units upward, so at coarse parcellations the limitation is dimensionality and not correlation. An investigator whose question is anatomical should specify the coarsest parcellation the question tolerates, before collection rather than after.

Reporting should name the equivalence class instead of the label. In our design, 58.6% of subregion pairs were exchangeable, in that substituting one for the other changed leave-one-out *R*^2^ by less than 0.002. Selecting one member of a redundant set and reporting it is a choice the data did not make, and the redundancy can be computed from the design alone, with no reference to any outcome.

An estimator that declines to name anything is behaving correctly here. The regularized horseshoe selected no subregion in most fits, and at the full sample with a 5% planted effect it selected something in 41.2% of repeats, with the selection containing a genuinely non-zero subregion in 94.2% of those. A procedure that always returns a maximum converts an unidentifiable design into a confident anatomical claim, which is how a literature arrives at two meta-analyses that disagree on sign.

### 4.7 Limitations

The whitened design cannot be acquired. Its purpose is decompositional, splitting the failure to recover into a part attributable to correlation and a part attributable to power, and only the granularity result translates into an instruction an investigator can follow.

Recovery depended on the planting scheme more than on collinearity. One true predictor gave 0.983, three of equal magnitude 0.692, three of decaying magnitude 0.511 and six of equal magnitude 0.300, against a paired collinearity cost of 0.158 at the same cell. The reported figure is one point on a calibration surface and not an estimate of what a real study would achieve. The direction of the dependence matters here: real effects are unlikely to be equal in magnitude, and the decaying pattern, the more plausible of the two, gave lower recovery than the configuration we report.

Resampling was performed within one cohort sharing scanner, protocol, preprocessing and population, so every agreement and recovery rate reported here bounds from above what independent replication would achieve. One dataset and one segmentation tool were used. Agreement between two automated subfield protocols across the adult lifespan is moderate, with correlations from r = 0.42 for the subiculum to r = 0.78 for CA1 (Samara et al., 2021), and reliability is not uniform across the labels we analysed: test-retest agreement is high for the molecular layer and dentate gyrus but lower for the parasubiculum (Brown et al., 2020) and the fimbria (Chiappiniello et al., 2021), three of which enter our design in both hemispheres. Both reliability studies used an earlier release of the segmentation module than the one analysed here. Measurement error and non-identifiabilitv are distinct problems that compound, and the specific numbers should be re-estimated before being transported to another pipeline.

Age absorbed 53% of outcome variance through the nuisance set. In a cross-sectional design that quantity does not separate age as a confounder from age as a mediator of a hippocampal pathway (Maxwell and Cole, 2007), and cross-sectional variance attributable to age is a poor proxy for longitudinal change (Lindenberger et al., 2011). Our observed null should be read as the absence of an age-independent association, consistent with longitudinal work in which change-change relationships appear only in specific subgroups, such as carriers of the APOE ε4 allele (Gorbach et al., 2017, 2020), and not as evidence that hippocampal volume is unrelated to cognitive ageing.

We predicted that divergent transitions would be more frequent in the real design than in the whitened one, giving a sampling-level signature of collinearity-induced multimodality. They were not. Median divergence rates were 0.0202 in both arms and scaled with sample size rather than with correlation. We report the prediction as tested and unsupported.

### 4.8 Conclusion

Subregional topography in the hippocampus is recoverable at a price, and at the effect sizes reported in cognitive ageing the price exceeds what any realistic study pays. Sample size buys the ability to detect that a structure matters well before it buys the ability to say which part of it does. The exchange rate between the two is set by the geometry of the parcellation, which is chosen, and not by the cohort, which is expensive.

## Information Sharing Statement

The Cam-CAN dataset analysed here is available to qualified researchers from the Cam-CAN data repository (https://camcan-archive.mrc-cbu.cam.ac.uk/dataaccess/) under its own data access agreement; the authors are not permitted to redistribute the raw imaging or cognitive data. FreeSurfer 8.2.0 is distributed at https://surfer.nmr.mgh.harvard.edu/. All analysis code written for this study, including the model definitions, the planted-effect experiment, the figure-generating scripts, the configuration files that fix every analytic decision, and the dated per-fit outputs and checkpoints of both planted-effect rounds, is archived at https://doi.org/10,5281/zenodo.21941295 under an open licence. Derived tables reporting every quantity in this article are included in that archive. Software versions for every dependency are listed in the Supplementary Material.

## Acknowledgments

We thank the participants who gave their time to the Cam-CAN protocol, without whose willingness to undergo extensive testing and scanning in the service of research this work would not exist.

Data collection and sharing for this project was provided by the Cambridge Centre for Ageing and Neuroscience (Cam-CAN). Cam-CAN funding was provided by the UK Biotechnology and Biological Sciences Research Council (grant number BB/H008217/1), together with support from the UK Medical Research Council and University of Cambridge, UK.

R.D. thanks the Coordenayao de Aperfeiyoamento de Pessoal de Nível Superior (CAPES) for its support. R.W. thanks CAPES and the Conselho Nacional de Desenvolvimento Cientifico e Tecnologico (CNPq) for their support.

## Statements and Declarations

### Ethics approval

The present work is a secondary analysis of an existing dataset and involved no new data collection. The Cam-CAN study was conducted in compliance with the Declaration of Helsinki and was approved by the local ethics committee, Cambridgeshire 2 Research Ethics Committee (reference 10/H0308/50). Written informed consent was obtained from all participants prior to their involvement.

### Author contributions

R.D.: conceptualisation, data curation, preprocessing, processing, methodology, software, formal analysis, investigation, visualisation, writing - original draft. R.W.: conceptualisation, methodology, formal analysis, writing - review and editing. Both authors read and approved the final manuscript.

### Data and code availability

See the Information Sharing Statement.

