## Supplementary Material for "Sample size buys detection, not localisation: an identifiability limit for hippocampal subfield morphometry"

This document contains Annex A, the expanded methodology, and Annex B, the supporting results. The two share a single page sequence and a single reference list but keep separate section, figure, table and equation series, so that a citation of Supplementary Table B4 or of equation A5.1 is unambiguous.

### Contents

**Annex A. Expanded methodology** **3**

A1 Notation 3

A2 Cohort assembly and label selection 3

A3 Nuisance adjustment and the Frisch–Waugh–Lovell equivalence 3

A4 Natural cubic spline basis for age 4

A5 Global and local decomposition 5

A6 Regularized horseshoe prior 5

A7 Quantities used for inference 6

A8 Exact leave-one-out $R^2$ for linear models 7

A9 Exchangeability and equivalence classes 7

A10 ZCA whitening 7

A11 Planted-effect construction 8

A12 Recovery metrics 8

A13 Saturation model and its bootstrap 8

A14 Sampling and diagnostics 9

A15 The segmentation atlas 9

A16 Computational environment 10

A17 Workflow 12

A18 Locked parameters 12

A19 Deviations from the analysis as first specified 14

**Annex B. Supporting results** **15**

B1 Per-subregion coefficients of the main model 15

B2 Robustness of the local null 16

B3 A prediction that failed: representative swapping 17

B4 Selection stability 18
B5 Projection predictive selection 20
B6 Multiverse 22
B7 Outcome specificity 24
B8 Prior sensitivity 25
B9 Age modelling 26
B10 Equivalence classes 26

### Annex A. Expanded methodology

This annex states every operation performed on the data in closed form, so that each step can be checked without reading the source code. It is written for a reader who wants to verify the arithmetic rather than accept it. Section numbering is independent of the main text.

#### A1 Notation

Let $n$ index participants and $p = 36$ the retained subregion labels. Write $\mathbf{V} \in \mathbb{R}^{n \times p}$ for the raw subregion volumes in mm$^3$, $\mathbf{v}_{\text{tot}} \in \mathbb{R}^n$ for whole hippocampal volume, $\mathbf{e} \in \mathbb{R}^n$ for estimated intracranial volume and $\mathbf{y} \in \mathbb{R}^n$ for the cognitive outcome. Vectors are column vectors, $\mathbf{1}_p$ is a vector of ones of length $p$, and $\mathcal{Z}(\cdot)$ denotes standardisation of each column to zero mean and unit variance with denominator $n - 1$:

$$\mathcal{Z}(\mathbf{u})_i = \frac{u_i - \bar{u}}{\text{sd}(\mathbf{u})}, \qquad \text{sd}(\mathbf{u})^2 = \frac{1}{n-1} \sum_{i=1}^{n} (u_i - \bar{u})^2. \tag{A1.1}$$

For a matrix $\mathbf{A}$ of full column rank, $\mathbf{P_A} = \mathbf{A}(\mathbf{A}^\top \mathbf{A})^{-1} \mathbf{A}^\top$ is the orthogonal projector onto its column space and $\mathbf{M_A} = \mathbf{I} - \mathbf{P_A}$ the residual maker.

#### A2 Cohort assembly and label selection

Subfield volumes were produced by the FreeSurfer 8.2.0 hippocampal subfield module, `segmentHA_T1.sh`, and exported with `asegstats2table`. From the exported table, columns corresponding to amygdala nuclei, to aggregate labels (whole hippocampus, head, body) and to the hippocampal fissure were removed bilaterally. No mask was constructed and no volume was recomputed; the operation was a selection of columns from the standard output. Estimated intracranial volume was taken from the same output.

The retained set is 18 labels per hemisphere: parasubiculum; presubiculum, subiculum, CA1, CA3, CA4, dentate gyrus and molecular layer, each split into anterior and posterior segments; fimbria; the hippocampus-amygdala transition area; and the hippocampal tail. CA2 is not a separate label in this atlas; the label denoted CA3 corresponds to the combined CA2 and CA3 field. Whole hippocampal volume, removed from the modelled set, was retained separately as the predictor of model M−1, so that the study reports what the global–local machinery buys over the simplest possible summary.

Participants with a missing value on the outcome or on any covariate were excluded, leaving $n = 638$ of 653. Missingness was assessed once, before any model was fitted.

#### A3 Nuisance adjustment and the Frisch–Waugh–Lovell equivalence

Volumes were transformed as $\log_{10}$, so that a multiplicative scaling of head size becomes an additive shift, and winsorised at $|z| = 4$ within column. Write $\mathbf{X}^{(0)} = \log_{10} \mathbf{V}$ after winsorisation.

The nuisance matrix is

$$\mathbf{Z} = \left[\ \mathbf{1} \mid \mathcal{Z}(\log_{10}\mathbf{e}) \mid \mathbf{S} \mid \mathbf{c}_{\text{sex}} \mid \mathcal{Z}(\mathbf{c}_{\text{edu}})\ \right], \tag{A3.1}$$

where $\mathbf{S} \in \mathbb{R}^{n\times 4}$ holds the natural cubic spline basis in age of Section A4 and $\mathbf{c}_{\text{sex}}$ is mean-centred. Both sides of the model were residualised on the same matrix,

$$\tilde{\mathbf{X}} = \mathbf{M_Z}\mathbf{X}^{(0)}, \qquad \tilde{\mathbf{y}} = \mathbf{M_Z}\mathbf{y}. \tag{A3.2}$$

By the Frisch–Waugh–Lovell theorem this is not a different model from entering $\mathbf{Z}$ alongside the predictors of interest. If $\boldsymbol{\beta}$ solves the full least-squares problem $\min\|\mathbf{y} - \mathbf{Z}\boldsymbol{\gamma} - \mathbf{X}^{(0)}\boldsymbol{\beta}\|^2$, then the same $\boldsymbol{\beta}$ solves $\min\|\tilde{\mathbf{y}} - \tilde{\mathbf{X}}\boldsymbol{\beta}\|^2$. The reason for residualising rather than including $\mathbf{Z}$ in the sampled model is that the shrinkage prior of Section A6 should act on the subregion contrasts alone; a covariate placed under that prior would be shrunk toward zero, which is not the intent for age or head size. The equivalence is exact for the least-squares problem and holds to the extent that the observation model is conditionally Gaussian; with the Student-$t$ likelihood actually used, residualisation is a first-order approximation, and this is the one place in the pipeline where the two formulations are not identical.

Nuisance adjustment absorbed a median of 0.419 of the variance of the subregion volumes (range 0.125 to 0.568) and 0.533 of the variance of the outcome.

#### A4 Natural cubic spline basis for age

Age enters through the truncated power basis for natural cubic splines of Hastie et al. (2009, Section 5.2.1). With knots $\xi_1 < \cdots < \xi_K$ placed at the quantiles $\{0.05, 0.275, 0.50, 0.725, 0.95\}$ of standardised age, so $K = 5$, define

$$d_k(x) = \frac{(x-\xi_k)_+^3 - (x-\xi_K)_+^3}{\xi_K - \xi_k}, \qquad k = 1, \ldots, K-1, \tag{A4.1}$$

where $(u)_+ = \max(u, 0)$. The basis used is

$$N_1(x) = x, \qquad N_{k+1}(x) = d_k(x) - d_{K-1}(x), \quad k = 1, \ldots, K-2, \tag{A4.2}$$

giving $K - 1 = 4$ columns beyond the intercept, each standardised. The construction forces the fitted function to be linear beyond $\xi_1$ and $\xi_K$, which is what prevents the erratic boundary behaviour of a high-order polynomial at the extremes of the age range. Linearity beyond the boundary knots was verified numerically by finite second differences.

The choice was compared against quadratic, linear and no age term by exact leave-one-out $R^2$ of the nuisance set on the outcome, computed as in Section A8. The spline and the quadratic were indistinguishable in this cohort (0.533 versus 0.537); the spline was retained because it had been specified in advance and because the comparison does not favour the quadratic.

#### A5 Global and local decomposition

Let $\mathbf{X}^*$ be $\tilde{\mathbf{X}}$ with columns standardised. The global factor is the standardised unweighted row mean,

$$\mathbf{g} = \mathcal{Z}\Big(\tfrac{1}{p}\,\mathbf{X}^*\mathbf{1}_p\Big)\,. \tag{A5.1}$$

An unweighted mean was used rather than the first principal component so that the factor does not depend on the sample: a principal component would be re-estimated in every subsample of the resampling analyses, and the local contrasts would then be defined against a moving target.

Local contrasts are obtained by removing the global factor and restandardising,

$$\mathbf{X}_{\text{loc}} = \mathcal{Z}\big(\mathbf{M}_{[\mathbf{1},\,\mathbf{g}]}\,\mathbf{X}^*\big)\,. \tag{A5.2}$$

By construction $\mathbf{X}_{\text{loc}}^\top\mathbf{g} = \mathbf{0}$, so the global coefficient and the local coefficients answer separate questions and are not competing for the same variance. Numerically the largest $|\mathbf{x}_{\text{loc},j}^\top\mathbf{g}|/n$ was $3.22 \times 10^{-16}$.

Equation (A5.2) removes the correlation between each local column and $\mathbf{g}$; it does not remove the correlation *among* the local columns, which after adjustment retained a first principal component accounting for 42% of variance, a condition number of 1,727 and a maximum variance inflation factor of 59.3. That residual structure is the object of the study.

#### A6 Regularized horseshoe prior

Local coefficients carry the regularized horseshoe of Piironen and Vehtari (2017), in the non-centred parameterisation. For $j = 1, \dots, p$,

$$\beta_j \mid \lambda_j, \tau, c \sim \mathcal{N}\Big(0,\ \tau^2\tilde{\lambda}_j^2\Big)\,, \qquad \tilde{\lambda}_j^2 = \frac{c^2\lambda_j^2}{c^2 + \tau^2\lambda_j^2}, \tag{A6.1}$$

$$\lambda_j \sim \mathcal{C}^+(0,1), \qquad c^2 \sim \text{Inv-Gamma}\Big(\tfrac{\nu_s}{2},\ \tfrac{\nu_s s^2}{2}\Big)\,, \tag{A6.2}$$

with slab degrees of freedom $\nu_s = 4$ and slab scale $s = 1$ on the standardised coefficient scale. The global scale is

$$\tau = \tau_{\text{raw}} \cdot \tau_0 \cdot \sigma, \qquad \tau_{\text{raw}} \sim \mathcal{C}^+(0,1), \qquad \tau_0 = \frac{p_0}{p - p_0} \cdot \frac{1}{\sqrt{n}}, \tag{A6.3}$$

where $p_0$ is the prior guess for the number of relevant predictors, set to 4. Equation (A6.3) is the calibration that makes the prior interpretable: a flat half-normal on $\tau$ would, at $p = 36$ and $n = 638$, correspond to a prior expectation of roughly 33 non-zero coefficients, which is not a sparsity assumption at all.

The shrinkage factor for coefficient $j$ and the effective number of non-zero coefficients are

$$\kappa_j = \frac{1}{1 + n\,\sigma^{-2}\,\tau^2\,\tilde{\lambda}_j^2}, \qquad m_{\text{eff}} = \sum_{j=1}^{p}(1 - \kappa_j)\,, \tag{A6.4}$$

with $\kappa_j = 1$ denoting complete shrinkage. Both were monitored: the posterior mean of $m_{\text{eff}}$ was

1.89 against the prior guess of four.

The main model is

$$y_i \sim t_\nu\left(\alpha + \beta_g\, g_i + \mathbf{x}_{\mathrm{loc},i}^\top \boldsymbol{\beta},\ \sigma\right), \qquad \nu = \nu_0 + 2, \quad \nu_0 \sim \mathrm{Exp}(1/11), \tag{A6.5}$$

with $\alpha \sim \mathcal{N}(0, 0.5^2)$, $\beta_g \sim \mathcal{N}(0, 1^2)$ entered unpenalised, and $\sigma \sim$ Half-Normal(1). The offset of two in the degrees of freedom keeps the variance finite.

Sensitivity to the prior was assessed by sweeping $p_0$ over $\{1, 2, 4, 8, 16\}$ and the slab scale over $\{0.5, 1, 2\}$, refitting the model at each value.

### A7 Quantities used for inference

#### A7.1 Bayesian $R^2$ and the local share

With $\boldsymbol{\mu}$ the linear predictor evaluated at a posterior draw,

$$R^2_{\mathrm{Bayes}} = \frac{\mathrm{var}(\boldsymbol{\mu})}{\mathrm{var}(\boldsymbol{\mu}) + \sigma^2}, \qquad r^2_{\mathrm{local}} = \frac{\mathrm{var}(\mathbf{X}_{\mathrm{loc}}\boldsymbol{\beta})}{\mathrm{var}(\boldsymbol{\mu}) + \sigma^2}, \tag{A7.1}$$

each computed per draw so that both have a posterior. The quantity $r^2_{\mathrm{local}}$ is the one on which the null is asserted: it is a single scalar computed inside one model and therefore does not inherit the identifiability problem that afflicts the individual coefficients.

The difference in Bayesian $R^2$ between M1 and M0 is formed from independent draws of the two posteriors. This is wider than a properly paired difference would be, which is conservative in the direction that matters for a claim of equivalence.

#### A7.2 Why a per-coefficient region of practical equivalence is not used

Under a region-of-practical-equivalence decision rule, the posterior mass of a parameter inside an interval around zero is treated as evidence for a practically null effect. The probability computed from a univariate marginal is conditional on the other parameters being independent of it. With predictors as correlated as these, that condition fails, and the marginal mass can be inflated or deflated relative to the joint. The per-coefficient mass is therefore reported in the coefficient table under a column name that marks it as invalid, and is never used to support a conclusion. Equivalence claims are restricted to $\beta_g$, $r^2_{\mathrm{local}}$ and $\Delta R^2$, each a single well-identified scalar, with half-widths 0.05, 0.01 and 0.01 respectively.

#### A7.3 Predictive comparison

Expected log pointwise predictive density was estimated by leave-one-out cross-validation with Pareto-smoothed importance sampling. For models $A$ and $B$ fitted to the same data,

$$\widehat{\Delta\mathrm{elpd}} = \sum_{i=1}^{n} \left(\widehat{\mathrm{elpd}}_i^A - \widehat{\mathrm{elpd}}_i^B\right), \qquad \mathrm{se} = \sqrt{n}\, \mathrm{sd}_i\left(\widehat{\mathrm{elpd}}_i^A - \widehat{\mathrm{elpd}}_i^B\right). \tag{A7.2}$$

The standard error is computed from the pointwise differences, which is the correct paired quantity, and not from the two marginal standard errors. A raw difference below four units was treated as small irrespective of its standard error; above four it was compared against the standard error. Pareto $k$ diagnostics were inspected for every model, with $k > 0.7$ flagged.

#### A8 Exact leave-one-out $R^2$ for linear models

Several diagnostics require an honest out-of-sample $R^2$ for an ordinary least-squares fit, computed thousands of times. For a design $\mathbf{A}$ with hat matrix $\mathbf{H} = \mathbf{P}_{\mathbf{A}}$ and residuals $\mathbf{r} = \mathbf{y} - \mathbf{A}\hat{\boldsymbol{\beta}}$, the leave-one-out residuals have the closed form

$$r_{(i)} = \frac{r_i}{1 - h_{ii}}, \qquad R^2_{\text{loo}} = 1 - \frac{\sum_i r^2_{(i)}}{\sum_i (y_i - \bar{y})^2}, \tag{A8.1}$$

which is exact and costs one decomposition. This is what makes the multiverse honest: with 36 predictors the in-sample incremental $R^2$ is positive by construction, and both versions were recorded so that the gap between them, which is overfitting, is visible.

#### A9 Exchangeability and equivalence classes

For each subregion $j$, let $R^2_{\text{loo}}(j)$ be the exact leave-one-out $R^2$ of the design $[\mathbf{1},\ \mathbf{g},\ \mathbf{x}_{\text{loc},j}]$. A pair $(j, k)$ is exchangeable when

$$\left|R^2_{\text{loo}}(j) - R^2_{\text{loo}}(k)\right| < \varepsilon, \qquad \varepsilon = 0.002. \tag{A9.1}$$

Equivalence classes were obtained by average-linkage hierarchical clustering on the dissimilarity $1 - |\text{corr}|$ between local columns, cut at $1 - 0.60$. Neither quantity uses the outcome beyond the single-predictor fit, and the clustering uses only the design, so both stand whether or not any effect exists.

#### A10 ZCA whitening

Let $\boldsymbol{\Sigma} = \text{cov}(\mathbf{X}_{\text{loc}})$ with eigendecomposition $\boldsymbol{\Sigma} = \mathbf{U}\boldsymbol{\Lambda}\mathbf{U}^\top$. The whitening transform used is

$$\mathbf{W}_{\text{ZCA}} = \mathbf{U}\,\boldsymbol{\Lambda}^{-1/2}\,\mathbf{U}^\top, \qquad \mathbf{X}_{\text{w}} = \mathcal{Z}\big((\mathbf{X}_{\text{loc}} - \bar{\mathbf{X}}_{\text{loc}})\,\mathbf{W}_{\text{ZCA}}\big)\,, \tag{A10.1}$$

with eigenvalues floored at $10^{-8}$. Among all matrices $\mathbf{W}$ satisfying $\mathbf{W}^\top\boldsymbol{\Sigma}\mathbf{W} = \mathbf{I}$, the ZCA choice minimises $\mathbb{E}\|\mathbf{X}\mathbf{W} - \mathbf{X}\|^2$ and is therefore the whitening transform closest to the identity. This matters for the comparison: principal-component whitening, $\mathbf{W} = \boldsymbol{\Lambda}^{-1/2}\mathbf{U}^\top$, would rotate the columns, so the $j$th whitened column would no longer correspond to the $j$th subregion and the two arms would be recovering different objects. Under ZCA the mean absolute correlation between each original column and its whitened counterpart was high, and the maximum off-diagonal correlation of the whitened design was of order $10^{-15}$.

#### A11 Planted-effect construction

Fix a subset $S \subset \{1, \dots, p\}$ with $|S| = k$, signs $\varsigma_j \in \{-1, +1\}$ and relative weights $w_j > 0$. Define the direction $\mathbf{d}$ by $d_j = \varsigma_j w_j$ for $j \in S$ and $d_j = 0$ otherwise. Let $\boldsymbol{\epsilon}$ be a noise vector standardised to zero mean and unit variance. The outcome is

$$\mathbf{y} = s\,\mathbf{X}\mathbf{d} + \boldsymbol{\epsilon}, \tag{A11.1}$$

and the scale $s$ is solved so that the planted predictors account for exactly a share $\rho$ of outcome variance:

$$\frac{s^2\,\mathrm{var}(\mathbf{X}\mathbf{d})}{s^2\,\mathrm{var}(\mathbf{X}\mathbf{d}) + 1} = \rho \qquad \Longrightarrow \qquad s = \sqrt{\frac{\rho}{(1-\rho)\,\mathrm{var}(\mathbf{X}\mathbf{d})}}. \tag{A11.2}$$

Equation (A11.2) is solved *separately in each arm*, because $\mathrm{var}(\mathbf{X}\mathbf{d})$ differs between the real and the whitened design for the same $\mathbf{d}$. Holding $s$ fixed instead would confound the comparison, since the arms would then carry different effect sizes. What is held identical across arms is the subsample, the columns, $S$, the signs, the weights and $\boldsymbol{\epsilon}$; what is solved per arm is only $s$.

Weights were $w_j = 1$ for the equal pattern and $w_j = 1/j$ for the decaying pattern, so that in the latter one predictor dominates and the remainder are progressively harder to detect.

#### A12 Recovery metrics

For a coefficient vector $\hat{\boldsymbol{\beta}}$ and the planted set $S$, with $\mathcal{T}_m$ the indices of the $m$ largest $|\hat{\beta}_j|$,

$$\text{top-1 is true} = \mathbb{1}\left[\arg\max_j |\hat{\beta}_j| \in S\right], \qquad \text{recall} = \frac{|S \cap \mathcal{T}_3|}{|S|}. \tag{A12.1}$$

For the lasso path the analogous quantities use the order in which predictors enter rather than coefficient magnitude. For the horseshoe, a subregion counts as *selected* when its 89% posterior interval excludes zero; the selection is *correct* when it contains at least one member of $S$. The difference between the proportion of fits that select anything and the proportion whose selection is correct is the false-localisation rate against a known truth.

#### A13 Saturation model and its bootstrap

Recovery as a function of sample size was summarised by

$$A(n) = a - b\,n^{-1/2}, \tag{A13.1}$$

fitted by nonlinear least squares with $a$ bounded in $[0, 1.05]$ and $b \geq -5$. The ceiling $a$ answers whether more participants would ever suffice; the rate $b$ answers whether each added participant buys the same amount in two designs. The sample size implied by a target $a^*$ is

$$n^* = \left(\frac{b}{a - a^*}\right)^2, \qquad a > a^*, \tag{A13.2}$$

and is infinite otherwise. When a fitted ceiling met the upper bound of 1.05, a companion fit with $a$ constrained to 1 was computed, and the two were reported as a range, since the unconstrained value is then a lower bound on the requirement.

Both parameters were bootstrapped by resampling *repeats* within each sample size and refitting, rather than by resampling the handful of grid points, which would measure scatter about the fitted line rather than uncertainty in the means. For contrasts between arms the same repeat indices were resampled in both arms before refitting each, preserving the pairing the experiment was built to exploit; this narrowed the interval on the difference in $b$ at the smaller planted effect from $[0.76, 3.18]$ under independent resampling to $[1.20, 2.79]$.

#### A14 Sampling and diagnostics

Posteriors were sampled with the No-U-Turn sampler. The observational models used 8,000 draws after 2,000 tuning iterations across 8 chains with a target acceptance probability of 0.9; the planted-effect experiment used 1,000 draws after 1,000 tuning iterations across 4 chains, with fits run in parallel and chains sequential within each fit. Convergence was monitored by $\hat{R}$ and effective sample size for every parameter, and divergent transitions were recorded per fit.

Across the 1,280 fits of the main experiment and the 1,320 of the closing round, no fit failed to complete. The median divergence rate was 0.0202 and was indistinguishable between the real and the whitened design, which is reported in the main text as a hypothesis tested and not supported. Recovery was recomputed on the subset of fits below a 1% divergence ceiling as a sensitivity analysis.

#### A15 The segmentation atlas

The hippocampal subfield volumes analysed here come from the probabilistic atlas of Iglesias et al. (2015), distributed with FreeSurfer and used here through release 8.2.0 (Fischl, 2012). The atlas was built from fifteen autopsy specimens scanned at approximately 0.13 mm isotropic resolution using custom hardware, a resolution roughly three orders of magnitude smaller in voxel volume than the 1 mm isotropic scans on which it is typically applied. Those specimens were manually labelled into thirteen hippocampal substructures under a protocol written for the purpose, and the resulting labels were combined with manual labels from an in vivo dataset covering the surrounding structures, using an atlas-building algorithm based on Bayesian inference. The released atlas is represented on a tetrahedral mesh with 18,417 vertices.

Segmentation proceeds by treating the atlas as a generative model of the image: the algorithm infers the deformation of the mesh and the intensity parameters jointly, which is what allows it to adapt to variations in contrast and resolution arising from different scanners and pulse sequences. It can therefore segment T1 images, T2 images or their combination. The original report validated the method on three public datasets, replicating findings in mild cognitive impairment based on high-resolution T2 data and discriminating Alzheimer's disease from elderly controls with 88% accuracy from standard resolution T1 scans alone.

Two properties of this construction bear directly on the present study. First, when the in vivo contrast is poor, the internal division of the hippocampus is determined largely by the prior rather than by the image, so neighbouring subfield volumes inherit a shared source of variation that is not participant-specific anatomy. Second, several boundaries the protocol must draw are not visible even in the ex vivo training data, so their placement is by convention. Both mechanisms push adjacent labels toward the redundancy quantified in the main text, and neither is remedied by increasing sample size. Independent test–retest work reports high reliability for the molecular layer, dentate gyrus and whole hippocampus, with intraclass correlations above 0.95, and the lowest reliability for the parasubiculum and the hippocampal fissure, with intraclass correlations between 0.78 and 0.89, mean volume differences above 5% and overlap below 70% (Brown et al., 2020). Of those two, only the parasubiculum is retained here, the fissure having been removed as a cerebrospinal fluid space; multicentre reproducibility work reports comparable limitations for the fimbria (Chiappiniello et al., 2021). Agreement between two automated protocols across the adult lifespan is moderate, with correlations from $r = 0.42$ for the subiculum to $r = 0.78$ for CA1 (Samara et al., 2021). Both reliability studies used FreeSurfer 6.0, so applying their conclusions to 8.2.0 assumes the module has not changed materially (Sämann et al., 2022).

#### A16 Computational environment

All analyses ran on a single workstation; no cluster or cloud resources were used, and no step required the graphics processor. Table A1 gives the machine and Table A2 the environment. The conda environment was built for this study and is archived with the code.

**Supplementary Table A1:** Workstation.

| Component | Specification |
|---|---|
| Operating system | Ubuntu 24.04.4 LTS (Noble Numbat) |
| Processor | AMD Ryzen 9 5950X, 16 cores, 32 threads |
| Motherboard | ASUS ROG Crosshair VIII Dark Hero (AMD X570) |
| Memory | 128 GB DDR4-3200, 4 × 32 GB |
| Storage, system | Samsung 990 Pro 2 TB NVMe, PCIe 4.0 |
| Storage, data | Seagate Exos X24 20 TB, SATA |
| Graphics | NVIDIA GeForce RTX 3090, 24 GB GDDR6X |

**Supplementary Table A2:** Software environment.

| Component | Version | Reference |
|---|---|---|
| FreeSurfer | 8.2.0 | Fischl (2012) |
| Hippocampal subfield atlas | as distributed | Iglesias et al. (2015) |
| conda | 26.3.2 | |
| PyMC | 6.0.1 | Abril-Pla et al. (2023) |
| ArviZ | 1.3.0 | Martin et al. (2026) |
| NumPy | 2.4.6 | Harris et al. (2020) |
| SciPy | 1.18.0 | Virtanen et al. (2020) |
| scikit-learn | 1.9.0 | Pedregosa et al. (2011) |
| pandas | 3.0.5 | |
| matplotlib | 3.8.4 | |
| seaborn | 0.13.2 | |

Sampling used the No-U-Turn sampler as implemented in PyMC, with chains run in parallel for the observational models and fits run in parallel with sequential chains for the planted-effect experiment, which is the arrangement that saturates 32 threads without oversubscribing them. Total sampling time for the 2,600 posterior fits of the two planted-effect rounds was a few hours. statsmodels 0.14.6 was present in the environment; no analysis reported here depends on it.

### A17 Workflow

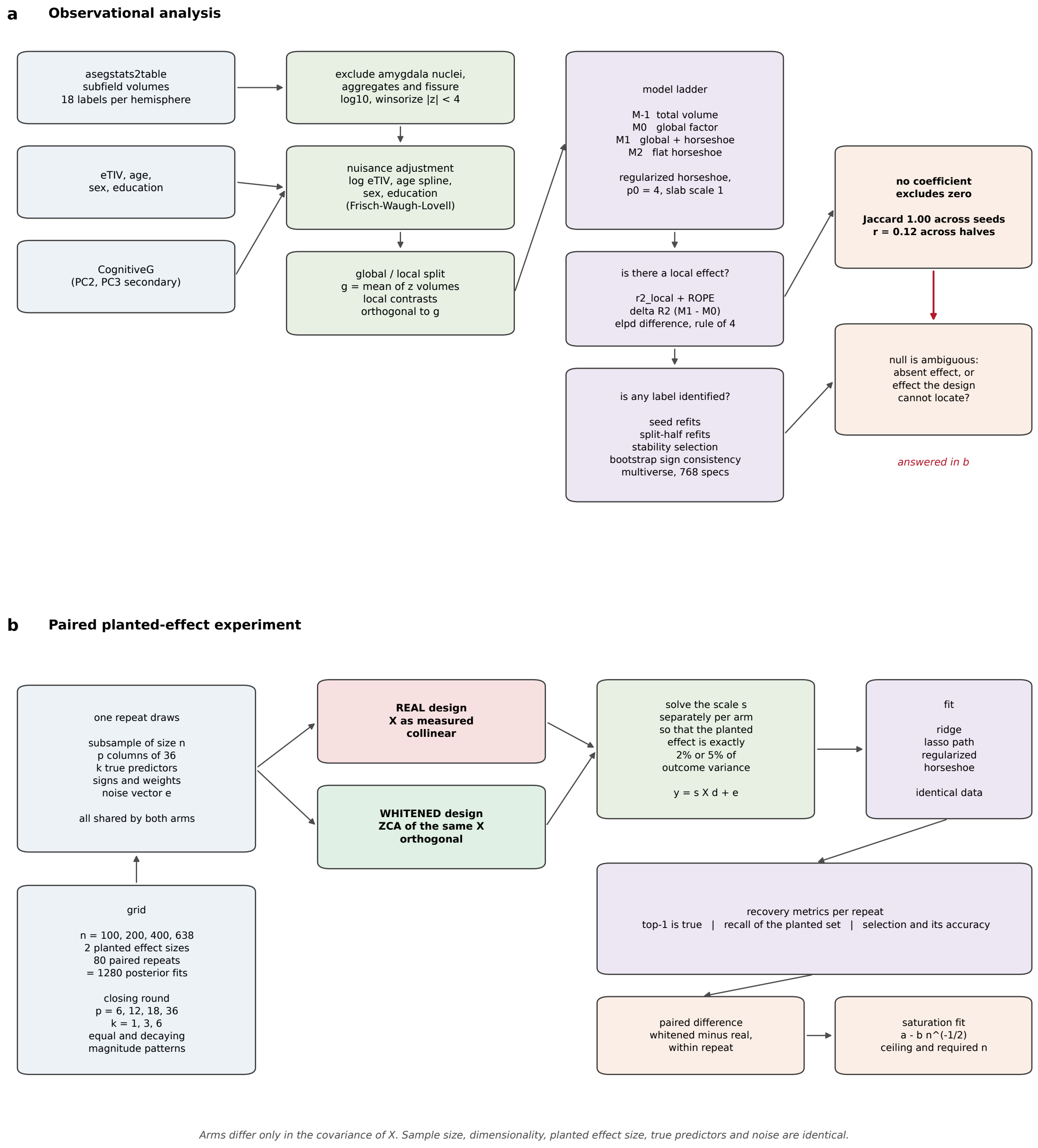


**Supplementary Figure A1:** Workflow. (a) The observational analysis, from label extraction to the three inferential quantities and the instability diagnostics, ending in an ambiguity the observed data cannot resolve. (b) The paired planted-effect experiment that resolves it. Within a repeat, the subsample, the columns, the true predictors, their signs and weights and the noise vector are shared by both arms; only the coefficient scale is solved separately, so that the planted effect accounts for exactly the target share of outcome variance in each. The arms therefore differ only in the covariance of the design.

### A18 Locked parameters

| Decision | Value | Basis |
|---|---|---|
| Volume transform | $\log_{10}$ | multiplicative scaling becomes additive |
| Winsorisation | $\lvert z\rvert = 4$ within column | one failed segmentation would dominate a 36-column correlation matrix |
| Head size | eTIV as covariate | proportion method biased when eTIV covaries with sex and age |
| Age | natural cubic spline, 5 quantile knots | nonlinear subcortical decline across the lifespan |
| Other covariates | sex, years of education | standard in this literature |
| Global factor | unweighted mean of standardised columns | does not move between subsamples, unlike a principal component |
| Local contrasts | residualised on $\mathbf{g}$, re-standardised | makes local and global questions separable |
| Shrinkage prior | regularized horseshoe | continuous analogue of spike-and-slab with calibrated sparsity |
| $p_0$ | 4 of 36 | prior guess for relevant predictors; swept over $\{1, 2, 4, 8, 16\}$ |
| Slab | $\nu_s = 4$, $s = 1$ | regularises the largest coefficients; swept over $s \in \{0.5, 1, 2\}$ |
| Likelihood | Student-$t$, $\nu = \nu_0 + 2$ | robustness to outlying observations |
| Interval mass | 89% equal-tailed | avoids the false precision of 95% with these effective sample sizes |
| ROPE, $\beta_g$ | $\pm 0.05$ | standardised units |
| ROPE, $r^2_{\text{local}}$, $\Delta R^2$ | $\pm 0.01$ | one percent of outcome variance |
| elpd rule | small if $\lvert\Delta\rvert < 4$ | Sivula et al. |
| Recovery criterion | 0.80 | threshold for calling a design identifiable |
| Planted effects | $k \in \{1, 3, 6\}$; equal and decaying | both varied because both were arbitrary |
| Planted effect sizes | 2% and 5% of outcome variance | generous relative to this literature |
| Whitening | ZCA | closest to the identity; preserves column identity |
| Exchangeability tolerance | 0.002 in $R^2_{\text{loo}}$ | below the resolution of the data |
| Clustering cut | $\lvert r\rvert \geq 0.60$, average linkage | |
| Saturation model | $a - b\,n^{-1/2}$, $a \in [0, 1.05]$ | classic root-$n$ form |
| Bootstrap | over repeats, pairing preserved | resampling grid points would understate uncertainty |

#### A19 Deviations from the analysis as first specified

Two departures are worth stating explicitly, because both would otherwise be invisible to a reader of the final text.

The identifiability grids were first computed with ridge regression as the workhorse, on the grounds that fitting a posterior in every cell of a four-way grid was not feasible. The estimator comparison later showed that ridge, which distributes coefficient mass across correlated predictors rather than selecting among them, recovers the planted set in 24.6% of repeats where the reported model reaches 69.2% and a lasso path reaches 76.7%. Every conclusion that had rested on the ridge grids was withdrawn, the parcellation gradient was recomputed with the reported model, and the estimator comparison was promoted from a robustness check to a result. The lesson is part of the paper's argument rather than an embarrassment to be buried: a study whose thesis is that analytic choices move findings by a factor of three should expect its own choices to do the same.

A hypothesis that divergent transitions would be more frequent in the collinear design than in the whitened one, and would therefore provide a sampling-level signature of collinearity-induced multimodality, was tested and not supported. Divergence rates were indistinguishable between arms and scaled with sample size. This is reported in the main text as a negative result rather than dropped.

### Annex B. Supporting results

This annex reports the analyses described in the Methods whose numerical results the main text summarises in a sentence or does not report at all. Every quantity here was produced by the same pipeline runs that produced the main results; nothing was refitted for this document.

#### B1 Per-subregion coefficients of the main model

The main text reports that no local coefficient was distinguishable from zero. Supplementary Table B1 gives all 36, so the claim can be checked rather than accepted. The largest posterior mean in absolute value is 0.0061 (Supplementary Table B1), against a global coefficient of 0.067, and the shrinkage factor exceeds 0.9 for every subregion. The final column carries the bivariate correlation with the outcome before the global factor is removed; comparing it with the posterior mean shows that no coefficient reverses sign relative to its own marginal association, which is what a suppression artefact would look like.

**Supplementary Table B1:** Posterior local coefficients of the main model, all 36 subregions. The shrinkage factor $\kappa$ is one under complete shrinkage. The final column is the bivariate correlation with the outcome before the global factor is removed, given so that any sign reversal is visible

| Subregion | Mean | SD | 89% lower | 89% upper | $\kappa$ | Marginal $r$ |
|---|---|---|---|---|---|---|
| L parasubiculum | -0.0003 | 0.0081 | -0.0089 | +0.0078 | 0.957 | 0.017 |
| L presubiculum (ant) | +0.0025 | 0.0118 | -0.0053 | +0.0180 | 0.945 | 0.090 |
| L presubiculum (post) | -0.0024 | 0.0118 | -0.0172 | +0.0055 | 0.945 | -0.009 |
| L subiculum (ant) | +0.0013 | 0.0094 | -0.0063 | +0.0129 | 0.952 | 0.081 |
| L subiculum (post) | -0.0009 | 0.0088 | -0.0109 | +0.0067 | 0.956 | 0.030 |
| L CA1 (ant) | +0.0010 | 0.0095 | -0.0066 | +0.0118 | 0.954 | 0.086 |
| L CA1 (post) | -0.0021 | 0.0112 | -0.0163 | +0.0058 | 0.946 | 0.012 |
| L CA3 (ant) | +0.0002 | 0.0101 | -0.0081 | +0.0099 | 0.953 | 0.064 |
| L CA3 (post) | -0.0002 | 0.0084 | -0.0088 | +0.0079 | 0.956 | 0.032 |
| L CA4 (ant) | +0.0043 | 0.0162 | -0.0047 | +0.0291 | 0.930 | 0.106 |
| L CA4 (post) | +0.0001 | 0.0085 | -0.0083 | +0.0084 | 0.958 | 0.050 |
| L DG (ant) | +0.0058 | 0.0203 | -0.0045 | +0.0390 | 0.919 | 0.110 |
| L DG (post) | -0.0002 | 0.0084 | -0.0088 | +0.0076 | 0.957 | 0.048 |
| L molecular layer (ant) | +0.0047 | 0.0172 | -0.0046 | +0.0321 | 0.927 | 0.109 |
| L molecular layer (post) | -0.0015 | 0.0102 | -0.0135 | +0.0061 | 0.950 | 0.035 |
| L fimbria | +0.0033 | 0.0132 | -0.0047 | +0.0228 | 0.940 | 0.083 |
| L HATA | +0.0023 | 0.0111 | -0.0055 | +0.0173 | 0.946 | 0.089 |
| L hippocampal tail | -0.0061 | 0.0192 | -0.0427 | +0.0041 | 0.915 | -0.019 |
| R parasubiculum | -0.0012 | 0.0091 | -0.0121 | +0.0063 | 0.954 | -0.011 |
| R presubiculum (ant) | +0.0018 | 0.0107 | -0.0059 | +0.0149 | 0.949 | 0.077 |
| R presubiculum (post) | -0.0005 | 0.0086 | -0.0097 | +0.0075 | 0.955 | 0.020 |
| R subiculum (ant) | +0.0004 | 0.0085 | -0.0075 | +0.0094 | 0.956 | 0.054 |
| R subiculum (post) | -0.0013 | 0.0094 | -0.0124 | +0.0061 | 0.953 | 0.025 |

| Subregion | Mean | SD | 89% lower | 89% upper | $\kappa$ | Marginal $r$ |
|---|---|---|---|---|---|---|
| R CA1 (ant) | -0.0014 | 0.0097 | -0.0129 | +0.0063 | 0.952 | 0.039 |
| R CA1 (post) | -0.0004 | 0.0083 | -0.0094 | +0.0075 | 0.957 | 0.032 |
| R CA3 (ant) | -0.0023 | 0.0117 | -0.0172 | +0.0055 | 0.945 | 0.019 |
| R CA3 (post) | +0.0030 | 0.0136 | -0.0051 | +0.0210 | 0.940 | 0.074 |
| R CA4 (ant) | -0.0025 | 0.0123 | -0.0173 | +0.0055 | 0.946 | 0.029 |
| R CA4 (post) | -0.0002 | 0.0086 | -0.0088 | +0.0081 | 0.954 | 0.041 |
| R DG (ant) | -0.0011 | 0.0097 | -0.0119 | +0.0068 | 0.952 | 0.041 |
| R DG (post) | -0.0004 | 0.0090 | -0.0095 | +0.0078 | 0.954 | 0.043 |
| R molecular layer (ant) | -0.0014 | 0.0099 | -0.0127 | +0.0063 | 0.952 | 0.049 |
| R molecular layer (post) | +0.0005 | 0.0088 | -0.0075 | +0.0093 | 0.955 | 0.055 |
| R fimbria | +0.0022 | 0.0111 | -0.0055 | +0.0165 | 0.947 | 0.074 |
| R HATA | -0.0005 | 0.0081 | -0.0100 | +0.0072 | 0.956 | 0.026 |
| R hippocampal tail | -0.0049 | 0.0171 | -0.0340 | +0.0045 | 0.925 | -0.022 |

#### B2 Robustness of the local null

Three quantities carry the null in the main text: the variance attributed to the local block within the main model, the difference in Bayesian $R^2$ against the model without it, and the difference in out-of-sample predictive fit. Supplementary Tables B2, B3 and B4 give them in full.

The null also survives two changes of specification that the Methods describe but the main text does not quantify. Removing the six labels with documented reliability limitations leaves 30 predictors and no interval excluding zero, with a largest absolute posterior mean of 0.0083. Replacing the local contrasts with centred log-ratio coordinates, which removes the geometric mean size exactly rather than removing an unweighted mean, gives 36 coordinates, again none excluding zero, with a largest absolute mean of 0.0058 (Supplementary Table B5). The two routes are not variations on a theme: one changes which labels enter, the other changes the geometry in which they are expressed. They agree with the main model and with each other.

**Supplementary Table B2:** Posterior Bayesian $R^2$ for each model in the ladder

| Model | Mean | SD |
|---|---|---|
| M-1 total volume | 0.0047 | 0.0052 |
| M0 global factor | 0.0068 | 0.0064 |
| M1 global + local | 0.0119 | 0.0091 |
| M2 flat horseshoe | 0.0070 | 0.0079 |

**Supplementary Table B3:** Pairwise differences in expected log pointwise predictive density against the global-only model, with standard errors computed from the pointwise differences

| Model | $\Delta$elpd | SE | Verdict |
|---|---|---|---|
| Mm1_total_only | -0.56 | 0.49 | small (\|diff\| < 4.0); prefer the simpler model |
| M1_global_plus_local | -0.39 | 0.59 | small (\|diff\| < 4.0); prefer the simpler model |
| M2_flat_horseshoe | -0.31 | 0.95 | small (\|diff\| < 4.0); prefer the simpler model |

**Supplementary Table B4:** Equivalence statements, restricted to the three well-identified scalars

| Quantity | Mean | 89% lower | 89% upper | P(in ROPE) | pd |
|---|---|---|---|---|---|
| beta_global | +0.0672 | +0.0044 | +0.1301 | 0.330 | 0.957 |
| r2_local | +0.0050 | +0.0000 | +0.0176 | 0.835 | 1.000 |
| delta_r2_M1_minus_M0 | +0.0051 | -0.0116 | +0.0240 | 0.640 | 0.684 |

**Supplementary Table B5:** Robustness of the local null across model specifications. No specification yields a single coefficient whose interval excludes zero

| Model | $p$ | Excluding zero | Max $\|\beta\|$ | Median $\|\beta\|$ |
|---|---|---|---|---|
| M1, main | 36 | 0 | 0.0061 | 0.0013 |
| M1r, reliable labels only | 30 | 0 | 0.0083 | 0.0018 |
| M1c, centred log-ratio | 36 | 0 | 0.0058 | 0.0014 |
| M2, flat horseshoe | 36 | 0 | 0.0102 | 0.0012 |

#### B3 A prediction that failed: representative swapping

We expected the posterior coefficients of anatomically adjacent subregions to anticorrelate. Under a sparsity-inducing prior with correlated predictors, the usual behaviour is that the model assigns weight to one member of a collinear group and withdraws it from the others, so that the joint posterior trades one label against its neighbour even when no single marginal is resolved. We looked for pairs whose posterior correlation fell below $-0.30$ and found none, in the main model or in the flat model without the global and local split (Supplementary Figure B1).

The explanation is visible in Annex A: the coefficients are shrunk too far for there to be anything to trade. With a posterior effective number of non-zero coefficients of 1.89 out of 36 and shrinkage factors above 0.9 throughout, the local block carries too little weight for the trade-off to appear. We record the prediction as tested and unsupported. Non-identifiability in this design presents as absent signal, not as signal that migrates between neighbours, which is a distinction worth having on record for anyone who expects the second symptom and concludes from its absence that the design is well behaved.

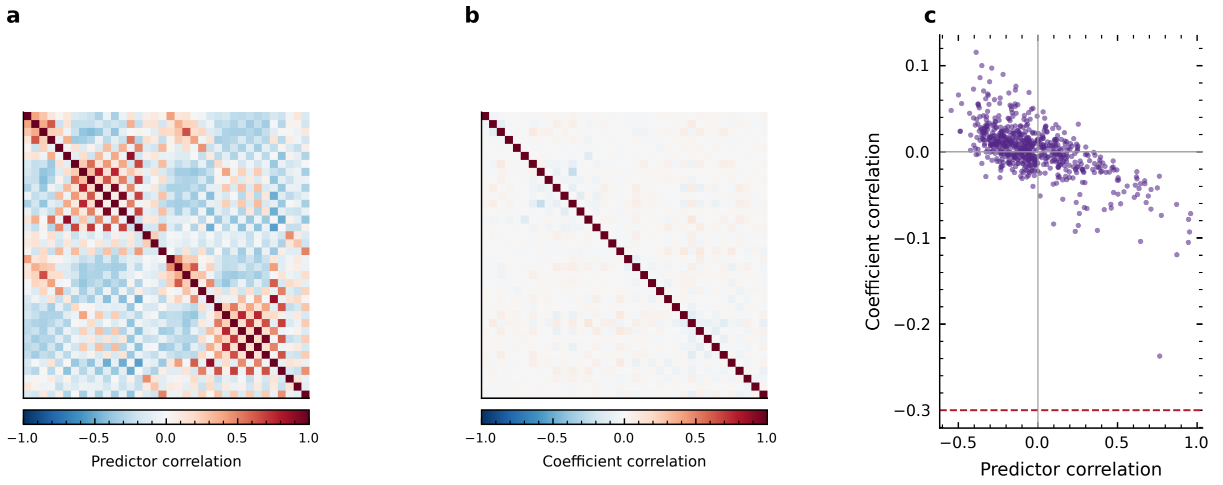


**Supplementary Figure B1:** Representative swapping between collinear neighbours, looked for and not found. Under a sparsity-inducing prior with correlated predictors the expected behaviour is that weight given to one member of a group is withdrawn from the others, which would appear as negative correlation between their posterior coefficients. (a) Correlation between the local contrasts entered in the model. (b) Correlation between their posterior coefficients, on the same colour scale. (c) The two plotted against each other for all 630 pairs, with the dashed line at the flagging threshold of -0.3; 0 pairs fall below it. The coefficients are shrunk too far for there to be anything to trade, so non-identifiability here presents as absent signal rather than as signal migrating between neighbours.

#### B4 Selection stability

Supplementary Table B6 gives the stability path for all 36 subregions, alongside two measures of sign consistency. No subregion is selected in a majority of subsamples: the highest selection frequency is 0.515 and the threshold of 0.60 is reached by none. Across bootstrap resamples the median sign consistency is 0.772, with 31 of 36 coefficients below the 0.90 criterion, so the sign of most coefficients is closer to a coin flip than to a finding.

Supplementary Tables B7 and B8 give the two resampling analyses whose contrast the main text uses as its central empirical observation. Refits differing only in the sampler seed agree perfectly, with a top-six overlap of 1.00 in all three pairwise comparisons. Refits on independent halves of the same cohort do not, with a mean coefficient correlation of +0.12 across six splits and individual splits ranging from −0.15 to +0.46.

**Supplementary Table B6:** Selection stability. Selection frequency is the proportion of 200 subsamples in which the subregion enters the first six of a lasso path; the two sign-consistency columns are computed within the selected subsamples and across bootstrap resamples of the full design respectively. No subregion reaches the 0.60 selection threshold

| Subregion | Selection frequency | Sign consistency, selection | Sign consistency, boots |
|---|---|---|---|
| R hippocampal tail | 0.515 | 0.961 | [illegible] |
| R CA3 (post) | 0.430 | 1.000 | [illegible] |
| L HATA | 0.400 | 1.000 | [illegible] |
| L hippocampal tail | 0.375 | 0.947 | [illegible] |
| L presubiculum (post) | 0.360 | 0.847 | [illegible] |
| R presubiculum (ant) | 0.320 | 1.000 | [illegible] |

| Subregion | Selection frequency | Sign consistency, selection | Sign consistency, boots[illegible] |
|---|---|---|---|
| L molecular layer (ant) | 0.315 | 1.000 | [illegible] |
| R fimbria | 0.270 | 0.981 | [illegible] |
| L fimbria | 0.265 | 1.000 | [illegible] |
| L CA4 (ant) | 0.265 | 1.000 | [illegible] |
| R CA4 (ant) | 0.230 | 0.891 | [illegible] |
| L DG (ant) | 0.220 | 1.000 | [illegible] |
| R subiculum (post) | 0.195 | 0.897 | [illegible] |
| L presubiculum (ant) | 0.175 | 0.971 | [illegible] |
| R CA3 (ant) | 0.155 | 0.774 | [illegible] |
| L parasubiculum | 0.135 | 0.519 | [illegible] |
| L CA1 (post) | 0.125 | 0.880 | [illegible] |
| R HATA | 0.115 | 0.565 | [illegible] |
| R CA1 (ant) | 0.110 | 0.818 | [illegible] |
| L subiculum (post) | 0.110 | 0.545 | [illegible] |
| R parasubiculum | 0.100 | 0.750 | [illegible] |
| R CA1 (post) | 0.080 | 0.562 | [illegible] |
| R DG (post) | 0.080 | 0.750 | [illegible] |
| R molecular layer (ant) | 0.080 | 0.875 | [illegible] |
| L CA3 (post) | 0.080 | 0.562 | [illegible] |
| L molecular layer (post) | 0.075 | 0.800 | [illegible] |
| R presubiculum (post) | 0.075 | 0.533 | [illegible] |
| L subiculum (ant) | 0.060 | 0.833 | [illegible] |
| R subiculum (ant) | 0.050 | 0.800 | [illegible] |
| R CA4 (post) | 0.045 | 0.667 | [illegible] |
| L CA4 (post) | 0.045 | 0.778 | [illegible] |
| L CA3 (ant) | 0.040 | 0.750 | [illegible] |
| R molecular layer (post) | 0.035 | 1.000 | [illegible] |
| L CA1 (ant) | 0.035 | 0.571 | [illegible] |
| L DG (post) | 0.030 | 0.667 | [illegible] |
| R DG (ant) | 0.005 | 1.000 | [illegible] |

**Supplementary Table B7:** Agreement between independent halves of the cohort, over six random splits with the full model refitted on each half

| Split | Top-6 overlap | Sign agreement | Coefficient correlation |
|---|---|---|---|
| 1 | 0.09 | 0.56 | +0.174 |
| 2 | 0.00 | 0.78 | +0.455 |
| 3 | 0.00 | 0.44 | +0.007 |
| 4 | 0.20 | 0.50 | -0.150 |
| 5 | 0.20 | 0.56 | +0.313 |
| 6 | 0.00 | 0.50 | -0.068 |

**Supplementary Table B8:** Overlap of the top-six selections between refits differing only in the sampler seed

| Seed pair | Top-6 overlap |
|---|---|
| 42 vs 7 | 1.00 |
| 42 vs 2718 | 1.00 |
| 7 vs 2718 | 1.00 |

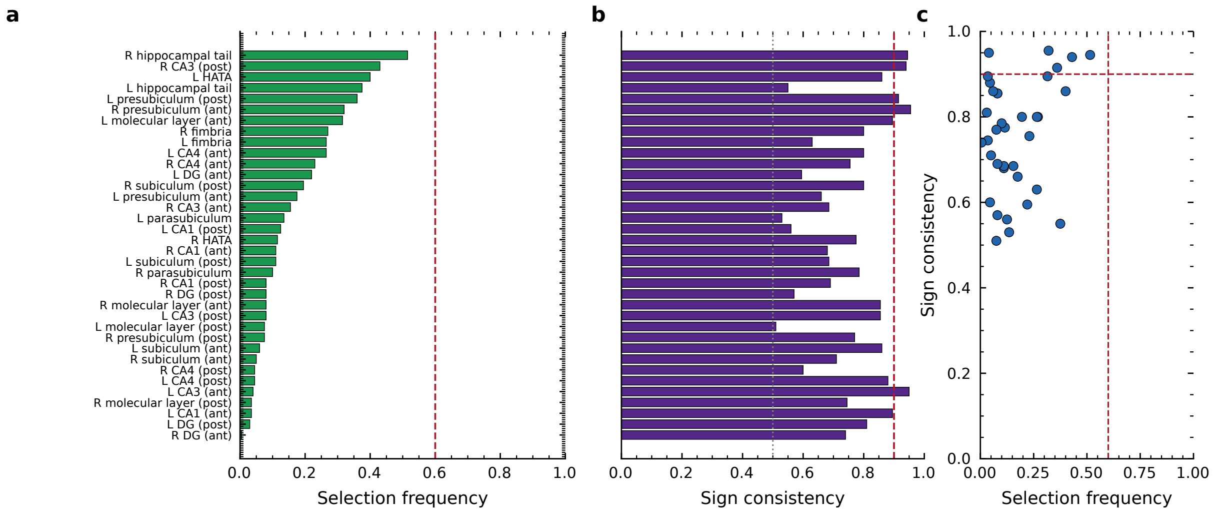


**Supplementary Figure B2:** Stability of subregional selection under resampling. (a) Selection frequency, the proportion of 200 subsamples of half the cohort in which each subregion enters the first six of a lasso path, with the dashed line at the selection threshold of 0.6; 0 subregions reach it. (b) Sign consistency, the proportion of bootstrap resamples of the full design in which each coefficient keeps its sign, with the dashed line at 0.9 and the dotted line at the value expected from a coin flip; the median is 0.772. (c) The two together. A subregion can be reported as a finding only if it sits in the upper right quadrant, selected often and with a stable sign, and none does.

#### B5 Projection predictive selection

The forward search adds local contrasts to a submodel that always contains the intercept and the global factor, so it decides only which contrasts earn a place. In sample, the best submodel gains 1.8 units of expected log pointwise predictive density over the global-only submodel, which is below the threshold of four at which a difference is treated as small (Supplementary Table B9). Under five-fold cross-validation, with the reference model refitted and the search repeated inside each training fold, the best gain is 0.08.

The more informative output is which subregion each fold selects first. Three distinct labels lead across five folds (Supplementary Table B10), so the identity of the leading subregion is a property of the training split.

**Supplementary Table B9:** Forward search of the projection predictive selection, in sample. The gain column is relative to a submodel containing only the intercept and the global factor

| Size | Added | elpd | Gain |
|---|---|---|---|
| 0 | (global only) | -902.9 | +0.00 |
| 1 | L hippocampal tail | -902.0 | +0.82 |
| 2 | L molecular layer (ant) | -901.5 | +1.41 |
| 3 | L DG (ant) | -901.4 | +1.47 |
| 4 | R hippocampal tail | -901.3 | +1.56 |
| 5 | R molecular layer (ant) | -901.2 | +1.68 |
| 6 | L HATA | -901.1 | +1.77 |
| 7 | R CA3 (post) | -901.1 | +1.79 |
| 8 | R fimbria | -900.9 | +1.97 |
| 9 | R DG (ant) | -900.8 | +2.11 |
| 10 | L presubiculum (ant) | -900.7 | +2.20 |
| 11 | L CA1 (post) | -900.6 | +2.22 |
| 12 | R subiculum (post) | -900.6 | +2.23 |

**Supplementary Table B10:** Subregion selected first by the forward search within each cross-validation fold

| Fold | First selected |
|---|---|
| 1 | L molecular layer (ant) |
| 2 | L hippocampal tail |
| 3 | L DG (ant) |
| 4 | L DG (ant) |
| 5 | L hippocampal tail |

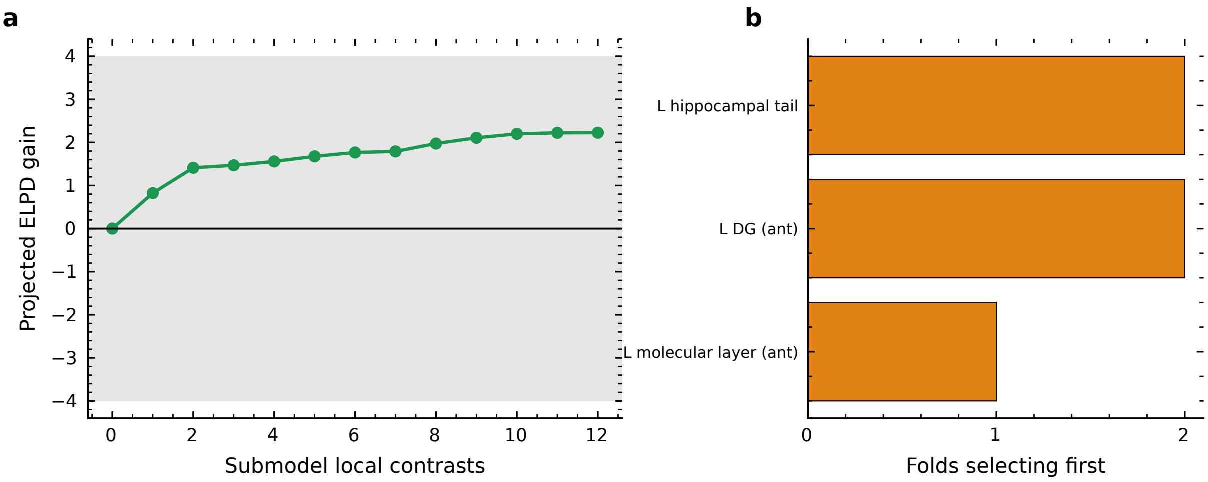


**Supplementary Figure B3:** Projection predictive variable selection. The forward search adds local contrasts to a submodel that always contains the intercept and the global factor, so it decides only which contrasts earn a place. (a) Projected gain in expected log pointwise predictive density, ELPD, of each submodel over the submodel containing the global factor alone, against the number of local contrasts it holds; the shaded band marks differences treated as small and the best gain is 2.2 units. The cross-validated version of this curve is reported in the text; its per-fold values were not retained in the archive. (b) Which subregion each fold of the cross-validation selects first; 3 distinct labels lead across 5 folds, so the identity of the leading subregion is a property of the training split rather than of the cohort. Subsegments are named anterior and posterior throughout.

#### B6 Multiverse

Across 768 specifications, crossing the method of intracranial volume adjustment, the age model, inclusion of sex and education, the label set, removal of the global factor, hemispheric treatment and winsorisation, the global coefficient stays stable in sign and magnitude while the identity of the largest local coefficient does not. Twelve different subregions take that place, the most frequent in fewer than a quarter of specifications (Supplementary Table B11). The incremental leave-one-out $R^2$ of the local block is negative in 576 of the 768 specifications, that is, in three quarters of them adding the local block makes out-of-sample prediction worse.

Supplementary Table B12 gives the vibration of effects by subregion. Coefficients that change sign across the specification space are not a minority.

**Supplementary Table B11:** Which subregion carries the largest local coefficient, across 768 analytic specifications

| Subregion | Specifications | Per cent |
|---|---|---|
| B DG (ant) | 231 | 30.1 |
| L molecular layer (ant) | 161 | 21.0 |
| R CA4 (ant) | 119 | 15.5 |
| B CA4 (post) | 77 | 10.0 |
| R DG (ant) | 50 | 6.5 |
| B molecular layer (post) | 40 | 5.2 |
| R molecular layer (post) | 29 | 3.8 |
| B molecular layer (ant) | 22 | 2.9 |
| L CA4 (post) | 14 | 1.8 |
| R molecular layer (ant) | 11 | 1.4 |
| B CA4 (ant) | 8 | 1.0 |
| B DG (post) | 6 | 0.8 |

**Supplementary Table B12:** Vibration of effects. Range of each subregion's coefficient across the specification space, collapsed to stem level so that the bilateral and separate-hemisphere specifications remain comparable

| Subregion | Mean | Minimum | Maximum | Sign consistency |
|---|---|---|---|---|
| fimbria | +0.075 | +0.018 | +0.241 | 0.500 |
| HATA | +0.013 | -0.023 | +0.045 | 0.573 |
| CA1 (post) | -0.033 | -0.260 | +0.026 | 0.711 |
| CA3 (post) | -0.054 | -0.331 | +0.044 | 0.767 |
| DG (post) | -0.076 | -0.479 | +0.352 | 0.798 |
| subiculum (ant) | -0.030 | -0.139 | +0.059 | 0.801 |
| CA4 (post) | +0.090 | -0.373 | +0.477 | 0.820 |
| hippocampal tail | -0.056 | -0.159 | +0.043 | 0.845 |
| subiculum (post) | -0.042 | -0.341 | +0.012 | 0.868 |

| Subregion | Mean | Minimum | Maximum | Sign consistency |
|---|---|---|---|---|
| presubiculum (ant) | +0.061 | -0.091 | +0.128 | 0.935 |
| molecular layer (ant) | +0.135 | -0.020 | +0.633 | 0.977 |
| parasubiculum | -0.028 | -0.072 | +0.009 | 0.984 |
| presubiculum (post) | -0.040 | -0.129 | +0.006 | 0.987 |
| CA4 (ant) | -0.202 | -0.933 | -0.017 | 1.000 |
| CA3 (ant) | -0.074 | -0.198 | -0.012 | 1.000 |
| CA1 (ant) | -0.120 | -0.447 | -0.014 | 1.000 |
| molecular layer (post) | +0.179 | +0.017 | +1.043 | 1.000 |
| DG (ant) | +0.297 | +0.080 | +1.225 | 1.000 |

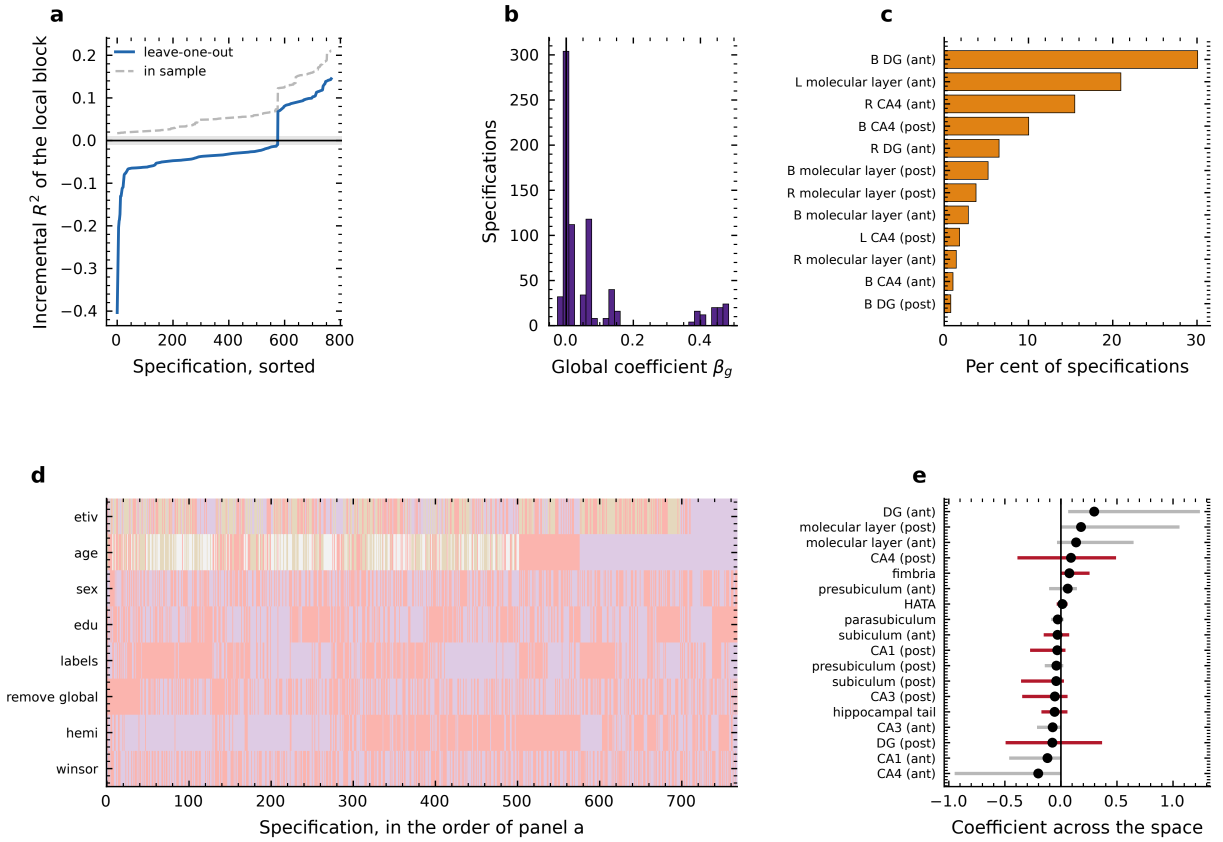


**Supplementary Figure B4:** Multiverse over 768 analytic specifications, crossing the method of intracranial volume adjustment, the age model, inclusion of sex and education, the label set, removal of the global factor, hemispheric treatment and winsorisation. (a) Incremental $R^2$ of the local block in every specification, sorted, computed by exact leave-one-out and, dashed, in sample; the gap between the two curves is overfitting, and the leave-one-out value is negative in 576 of 768 specifications. (b) Distribution of the global coefficient, which is stable in sign and magnitude throughout. (c) Which subregion carries the largest local coefficient, as a percentage of specifications; 12 different labels take that position. (d) The analytic decisions behind each specification, in the order of panel a, one row per decision. (e) Range of each subregion's coefficient across the whole space with its mean marked; 9 of 18 change sign somewhere in the space and are drawn in red.

#### B7 Outcome specificity

A multivariate model fitted the three cognitive components jointly with a full residual covariance. The global coefficient is +0.075 for the general factor, with an 89% interval of [0.010, 0.139], against +0.022 and +0.024 for the second and third components, whose intervals include zero (Supplementary Table B13).

The contrasts do not resolve. The difference between the general factor and the second component is +0.053 with an interval of [−0.031, 0.136], and against the third component +0.050 with [−0.036, 0.138]; both place between 44% and 47% of their posterior mass inside the region of practical equivalence. The association is therefore not demonstrably specific to general cognitive ability. Given that the association itself accounts for under one per cent of outcome variance, this is the expected result and not an additional finding, but it should be reported rather than left implicit.

**Supplementary Table B13:** Multivariate model over the three cognitive components. The first three rows are the global coefficient for each outcome; the last two are the contrasts that formalise the specificity claim

| Quantity | Mean | 89% lower | 89% upper | P(in ROPE) | pd |
|---|---|---|---|---|---|
| CognitiveG | +0.075 | +0.010 | +0.139 | 0.265 | 0.967 |
| CognitivePC2 | +0.022 | -0.042 | +0.085 | 0.729 | 0.707 |
| CognitivePC3 | +0.024 | -0.040 | +0.089 | 0.720 | 0.734 |
| CognitiveG vs CognitivePC2 | +0.053 | -0.031 | +0.136 | 0.447 | 0.851 |
| CognitiveG vs CognitivePC3 | +0.050 | -0.036 | +0.138 | 0.467 | 0.810 |

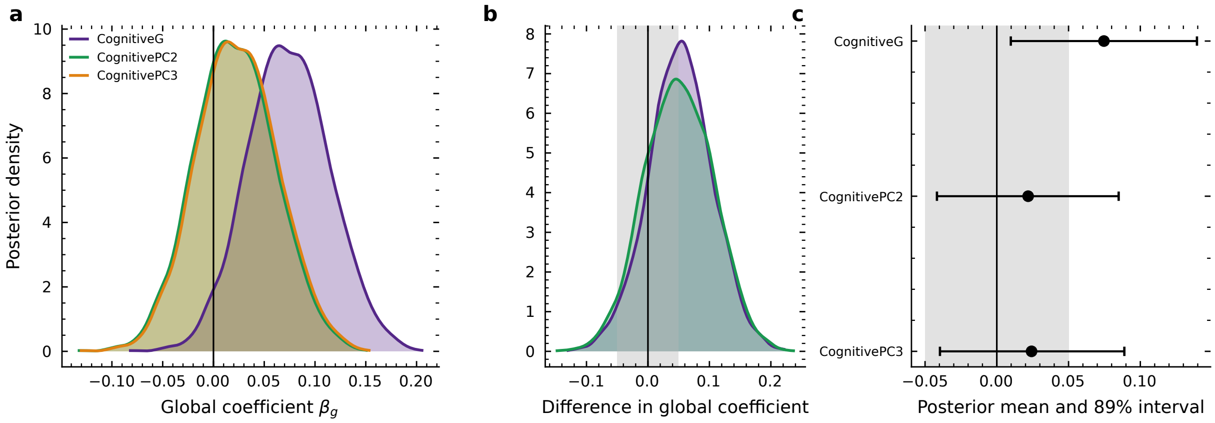


**Supplementary Figure B5:** Outcome specificity, from a multivariate model fitting the three cognitive components jointly with a full residual covariance. (a) Posterior of the global hippocampal coefficient for each component. (b) Posterior of the difference between the general factor and each secondary component, with the region of practical equivalence shaded: CognitiveG vs CognitivePC2 +0.053 [-0.031, +0.136]; CognitiveG vs CognitivePC3 +0.050 [-0.036, +0.138]. Both intervals include zero, so the association is not demonstrably specific to general cognitive ability. (c) The same quantities as posterior means with 89 per cent intervals, for the three outcomes. Given that the association accounts for under one per cent of outcome variance, this is the expected result rather than an additional finding.

#### B8 Prior sensitivity

The prior guess for the number of relevant predictors was swept from 1 to 16 and the slab scale from 0.5 to 2, refitting the model at each value (Supplementary Table B14). The global coefficient is unmoved, varying between 0.066 and 0.068 across the entire sweep. The variance carried by the local block rises with the prior guess, from 0.0024 at $p_0 = 1$ to 0.0074 at $p_0 = 16$, and remains below the equivalence threshold of 0.01 at every setting including the most permissive.

This addresses the obvious objection to a null obtained under a sparsity-inducing prior, which is that the prior produced it. A prior four times more permissive than the one adopted does not lift the local block above the threshold, and the posterior effective number of non-zero coefficients reaches only 2.88 when the prior expects 16.

**Supplementary Table B14:** Prior sensitivity. The global coefficient is unmoved across the whole sweep; the variance carried by the local block rises with the prior guess for the number of relevant predictors but remains an order of magnitude below the equivalence threshold

| **Swept** | **Value** | $m_{\text{eff}}$ | $r^2_{\text{local}}$ | $\beta_g$ |
|---|---|---|---|---|
| p0 | 1.0 | 0.88 | 0.0024 | 0.0667 |
| p0 | 2.0 | 1.29 | 0.0035 | 0.0678 |
| p0 | 4.0 | 1.86 | 0.0050 | 0.0677 |
| p0 | 8.0 | 2.42 | 0.0063 | 0.0674 |
| p0 | 16.0 | 2.88 | 0.0074 | 0.0661 |
| slab_scale | 0.5 | 1.96 | 0.0052 | 0.0668 |
| slab_scale | 1.0 | 1.86 | 0.0050 | 0.0677 |
| slab_scale | 2.0 | 1.88 | 0.0049 | 0.0674 |

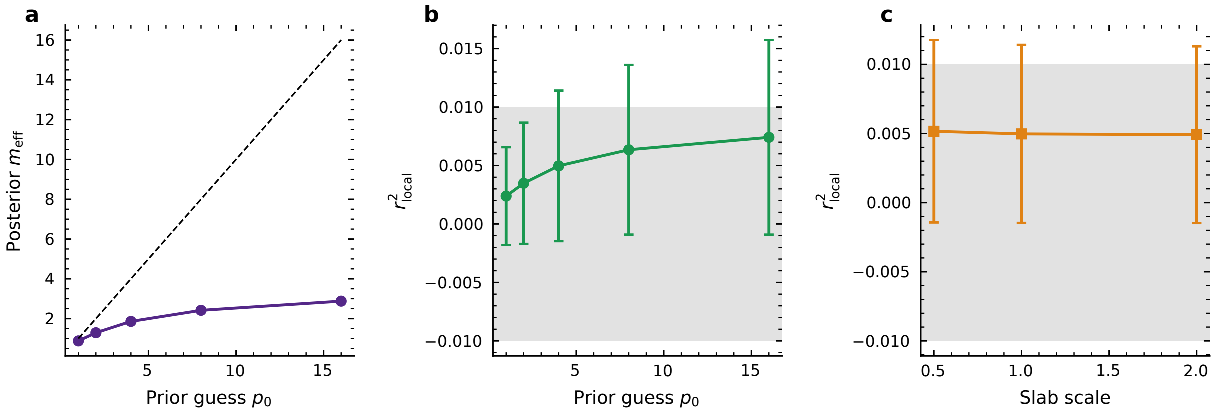


**Supplementary Figure B6:** Sensitivity of the result to the sparsity prior, which is the obvious objection to a null obtained under shrinkage. (a) Posterior effective number of non-zero coefficients against the prior guess $p\_0$, with the dashed identity line; the posterior never follows the prior upward, reaching 2.88 when the prior expects 16. (b) Variance carried by the local block as $p\_0$ is swept, with the region of practical equivalence shaded. (c) The same quantity as the slab scale is swept. Across the whole sweep the global coefficient moves only between 0.066 and 0.068, and the local block stays below the equivalence threshold at every setting, including the most permissive one, which expects four times as many relevant predictors as the reported analysis.

#### B9 Age modelling

Age was specified in advance as a natural cubic spline with five quantile knots. Supplementary Table B15 compares that choice against a quadratic, a linear term and no age term, by exact leave-one-out $R^2$ of the nuisance set on the outcome.

The spline and the quadratic are indistinguishable in this cohort, at 0.5333 and 0.5368, and the linear term is only slightly worse at 0.5167. Omitting age altogether drops the fit to 0.1940. The spline was retained because it had been specified before the comparison was made and because the comparison does not favour the quadratic; the choice does not affect any conclusion, since the same nuisance set is used for every model in the ladder and therefore cannot favour one over another.

**Supplementary Table B15:** Exact leave-one-out $R^2$ of the nuisance set on the outcome under each age specification

| Age model | Columns | Leave-one-out $R^2$ |
|---|---|---|
| spline | 8 | 0.5333 |
| quadratic | 6 | 0.5368 |
| linear | 5 | 0.5167 |
| none | 4 | 0.1940 |

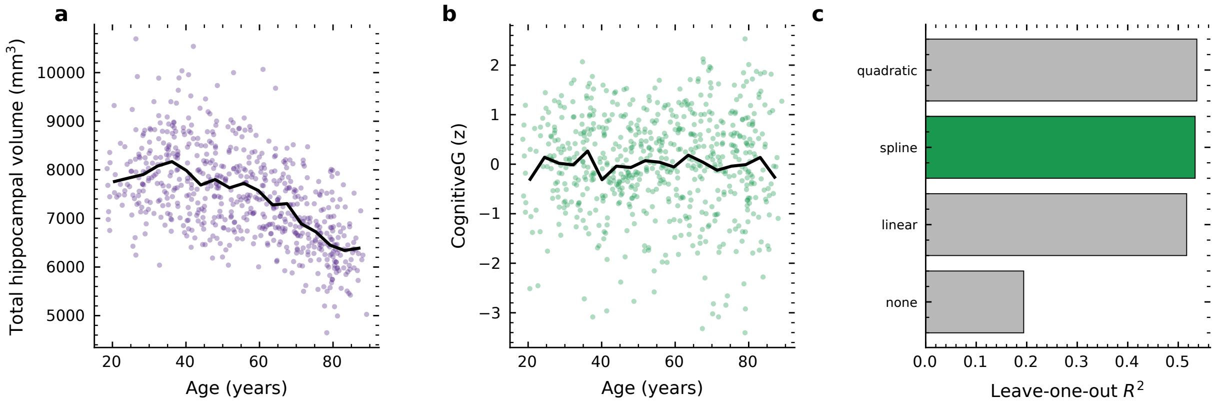


**Supplementary Figure B7:** Age, and how much of the outcome it absorbs. (a) Total hippocampal volume against age across the adult lifespan, with a running mean; the decline is not linear, which is why age enters as a spline. (b) The cognitive outcome against age, on the same axis. (c) Exact leave-one-out $R^2$ of the nuisance set on the outcome, that is, how much of CognitiveG the covariates predict out of sample, under each age specification, with the specification used in the reported analysis in green. The natural cubic spline reaches 0.5333 against 0.5368 for the best alternative, so the two are indistinguishable in this cohort; the spline was retained because it had been specified before the comparison was made. The same nuisance set enters every model in the ladder, so this choice cannot favour one over another.

#### B10 Equivalence classes

Supplementary Table B16 lists the equivalence classes by name. The grouping is computed from the design alone, without reference to any outcome, so it holds whether or not a subfield effect exists and can be computed by anyone with the same segmentation before they collect an outcome at all. It is the form in which we would recommend subfield results be reported: a class that cannot be separated by the data is the unit the data support, and naming one member of it is a choice the data did not make.

**Supplementary Table B16:** Equivalence classes of subregions, obtained by average-linkage clustering on one minus the absolute correlation between local contrasts, cut at 0.60. The grouping uses the design alone and no outcome

| Class | Size | Members |
|---|---|---|
| 1 | 4 | L CA3 (post), L CA4 (post), L DG (post), L molecular layer (post) |
| 2 | 1 | L CA1 (post) |
| 3 | 2 | R CA1 (ant), R molecular layer (ant) |
| 4 | 2 | R presubiculum (ant), R subiculum (ant) |
| 5 | 1 | L parasubiculum |
| 6 | 1 | R parasubiculum |
| 7 | 1 | L subiculum (post) |
| 8 | 1 | R subiculum (post) |
| 9 | 3 | R CA3 (ant), R CA4 (ant), R DG (ant) |
| 10 | 2 | L presubiculum (post), R presubiculum (post) |
| 11 | 2 | L presubiculum (ant), L subiculum (ant) |
| 12 | 4 | R CA3 (post), R CA4 (post), R DG (post), R molecular layer (post) |
| 13 | 1 | R CA1 (post) |
| 14 | 1 | L hippocampal tail |
| 15 | 1 | R hippocampal tail |
| 16 | 2 | L CA1 (ant), L molecular layer (ant) |
| 17 | 3 | L CA3 (ant), L CA4 (ant), L DG (ant) |
| 18 | 1 | L fimbria |
| 19 | 1 | R fimbria |
| 20 | 1 | L HATA |
| 21 | 1 | R HATA |
